# G-quadruplex structures regulate long-range transcriptional reprogramming to promote drug resistance in ovarian cancer

**DOI:** 10.1101/2024.06.24.600010

**Authors:** Jenna Robinson, Gem Flint, Ian Garner, Silvia Galli, Thomas E. Maher, Marina K. Kuimova, Ramon Vilar, Iain A. McNeish, Robert Brown, Hector Keun, Marco Di Antonio

**Affiliations:** Imperial College London, Department of Chemistry, Molecular Sciences Research Hub, 82 Wood Lane, London W12 0BZ, UK; Imperial College London, Institute of Chemical Biology, Molecular Sciences Research Hub, 82 Wood Lane, London W12 0BZ, UK; Department of Pediatric Oncology, Dana Farber Cancer Institute, Boston, MA 02215, USA; Division of Cancer, Department of Surgery and Cancer, Imperial College London, London W12 0NN, UK; The Francis Crick Institute, 1 Midland Road, London NW1 1AT, UK

**Author notes:** These authors contributed equally to this work (Jenna Robinson, Gem Flint). To whom correspondence should be addressed.; (MDA).

## Abstract

Epigenetic evolution is a common mechanism used by cancer cells to evade the therapeutic effects of drug treatment. In ovarian cancers, epigenetically-driven resistance may be responsible for a large number of late-stage patient deaths. Here, we describe the first investigation into the role of G-quadruplex (G4) DNA secondary structures in mediating epigenetic regulation in drug-resistant ovarian cancer cells. Through genome-wide mapping of G4s in paired drug-sensitive and drug-resistant cell lines, we find that increased G4 formation is associated with significant increase in gene expression, with high enrichment in signalling pathways previously established to promote drug-resistant states. However, in contrast to previous studies, the expression-enhancing effects of G4s were not found at gene promoters, but intergenic and intronic regions, indicating that G4s promote long-range transcriptional regulation in drug-resistant cells. Furthermore, we discovered that clusters of G4s (super-G4s) are associated with particularly high levels of transcriptional enhancement that surpass the effects of super-enhancers, which act as well established regulatory sites in many cancers. Finally, we demonstrate that targeting G4s with small molecules results in significant down-regulation of pathways associated with drug-resistance, which results in resensitisation of resistant cells to chemotherapy agents. These findings indicate that G4 structures are critical for the epigenetic regulatory networks of drug-resistant cells and may represent a promising target to treat drug-tolerant ovarian cancer.

## Introduction

Drug-resistance is the greatest challenge associated with the treatment of ovarian cancer patients, with 80% of late-stage patients acquiring some form of drug resistance during the course of their treatment.^1^ Often drug-resistant states are attained via epigenetic adaption, including altered expression of DNA damage repair pathways and drug transporter proteins that allow cells to combat the effects of DNA damaging drugs.^1,2^ Much effort has been directed at understanding the molecular mechanism that drive such epigenetic changes, with a large focus being placed on histone modifications as well as DNA methylation states.^3,4^ However, very limited work has considered the non-canonical DNA structures that form within cells, and how deviations from the common duplex structure of DNA may influence epigenetic processes in drug-resistant cells.

G-quadruplexes (G4s) are amongst the most studied DNA secondary structures which form when guanine bases Hoogsteen base pair to form stacks of G-quartets.^5,6^ The biological relevance of G4s has been established by means of multiple sequencing and detection methods including *in vitro* sequencing approaches such as G4-seq,^7^ as well as cellular sequencing methods that utilise G4-specific antibodies such as BG4 ChIP-seq and BG4 CUT&Tag.^8,9^ Cellular detection of G4s has led to their association with multiple diseases, notably revealing a global enrichment of G4s in cancer cells, which has now been extensively validated by multiple independent G4 detection methods.^10–14^

The most studied genomic location associated with G4 formation is gene promoters, including those of known oncogenes such as *c-MYC, KRAS* and *BCL-2*.^15,16^ Furthermore, endogenous G4 formation at gene promoters has been linked to active gene expression in multiple cancer cell lines and patient-derived xenograft models including in keratinocytes, liposarcomas and breast cancer cells.^11,17–20^ There are a number of mechanisms that may explain such associations, including the notion that G4s can act as binding platforms for various transcription factors and regulatory proteins and may also influence chromatin architecture.^21,22^ The impact that G4s have on the epigenetic state of cells may therefore be utilised by cancer cells to rapidly adapt their gene expression profile to external stressors. This hypothesis is supported by a recent study demonstrating that the acquisition of temozolomide-resistance in glioblastomas is associated with a decrease in G4 prevalence.^23^

In the case of drug-resistant ovarian cancer, recent work has highlighted the critical importance of transcriptional enhancer regions found outside of promoters for long-range regulation of gene expression.^24–26^ Furthermore, multiple studies have revealed that clusters of enhancer sites known as super-enhancers are key orchestrators of the drug response in ovarian cancer,^25,26^ repeatedly facilitating larger changes of gene expression than those obtained by modification of individual promoters. Importantly, G4s too have been linked to distal control of gene expression through their enrichment at the boundaries of loops linking enhancer and promoters as well as their direct recruitment of loop-mediating proteins such as YY1 and CTCF.^27–30^ G4s can additionally form intermolecularly between strands of DNA which results in distinct recognition by chromatin remodellers such as CSB,^31^ and phase-separation events,^32^ which have both been previously linked to increased regulatory protein recruitment across long distances.^33,34^ We therefore hypothesised that G4 formation may also be used by ovarian cancer cells to epigenetically drive drug resistance, either proximally at promoters or distally via long-range mechanisms.

To investigate this hypothesis, we undertook a multi-layered genomics approach, sequencing paired drug-sensitive and drug-resistant ovarian cancer cell lines. Specifically, we utilised the G4-specific antibody BG4 to obtain G4 maps, firstly, in the established drug-sensitive and drug-resistant ovarian cancer cell lines PEO1 and PEO4,^35^ in addition to RNA-sequencing, chromatin accessibility mapping (ATAC-seq) and enhancer profiling (H3K27ac CUT&Tag). From this data we found that the association of G4 formation with gene expression was tightly linked to their genomic context. Surprisingly, the gain of new G4 sites in PEO4 cells only significantly impacted gene expression when detected outside of promoter regions, which was not observed in any of previous genomic studies on G4s. This observation was further validated in additional paired ovarian cancer lines (PEA1/PEA2). Furthermore, small-molecule targeting of G4s was leveraged to perturb G4-homeostasis and epigenetically resensitise drug-resistant PEO4 cells to cisplatin. RNA-sequencing within resistant cells treated with the G4-ligand PDS revealed downregulation of key genes associated with cisplatin resistance that contained a G4-peak in a non-promoter region, explaining the observed re-sensitisation to cisplatin treatment of PDS treated PEO4 cells. Our results indicate that G4s are important for establishing transcriptional networks leveraged by drug-resistant ovarian cancer cells and that disrupting the G4 landscape of drug resistant cells may be a viable therapeutic approach to counteract the transcriptional rewiring that underpins drug-tolerance.

## Results

### G4s at non-promoter regions are associated with elevated gene expression in drug-resistant cells

To investigate how changes in G4 distribution correlate with epigenetic reprogramming observed in drug-resistant ovarian cancer cells, we set out to map G4s within the chromatin of a pair of high-grade serous ovarian carcinoma cell lines, PEO1 and PEO4.^35^ Importantly, PEO1 and PEO4 cells were established from a single patient before and after (respectively) she developed resistance to a mixture of chemotherapies, including cisplatin, chlorambucil and 5-fluorouracil.^35^ G4-mapping was initially performed via ChIP-seq with the G4-selective antibody BG4.^11,36^ Initially, antibody selectivity was confirmed by conducting ChIP-qPCR with primers targeting validated G4-forming sequences taken from the literature.^37^ This yielded an 8-fold enrichment for G4 sites after BG4 immunoprecipitation compared to non-G4 regions (Figure 1A), which was encouragingly beyond the 5-fold enrichment threshold recommended for successful G4 immunoprecipitation.^37^ BG4 ChIP-seq was thus extended to both PEO1 and PEO4 and peaks present in at least two of the three biological replicates were taken forward for further analysis, in agreement with the literature.^37^ To validate selective enrichment of G4 structures, we used MEME suite to extract enriched sequence motifs within the top 1,000 G4 peaks. This revealed the most enriched motif was a G-rich sequence in ChIP-seq peaks found in both cell lines (Figure 1B, S1), further indicating the generation of reliable G4 maps in both PEO1 and PEO4 cells.

**Figure 1.**
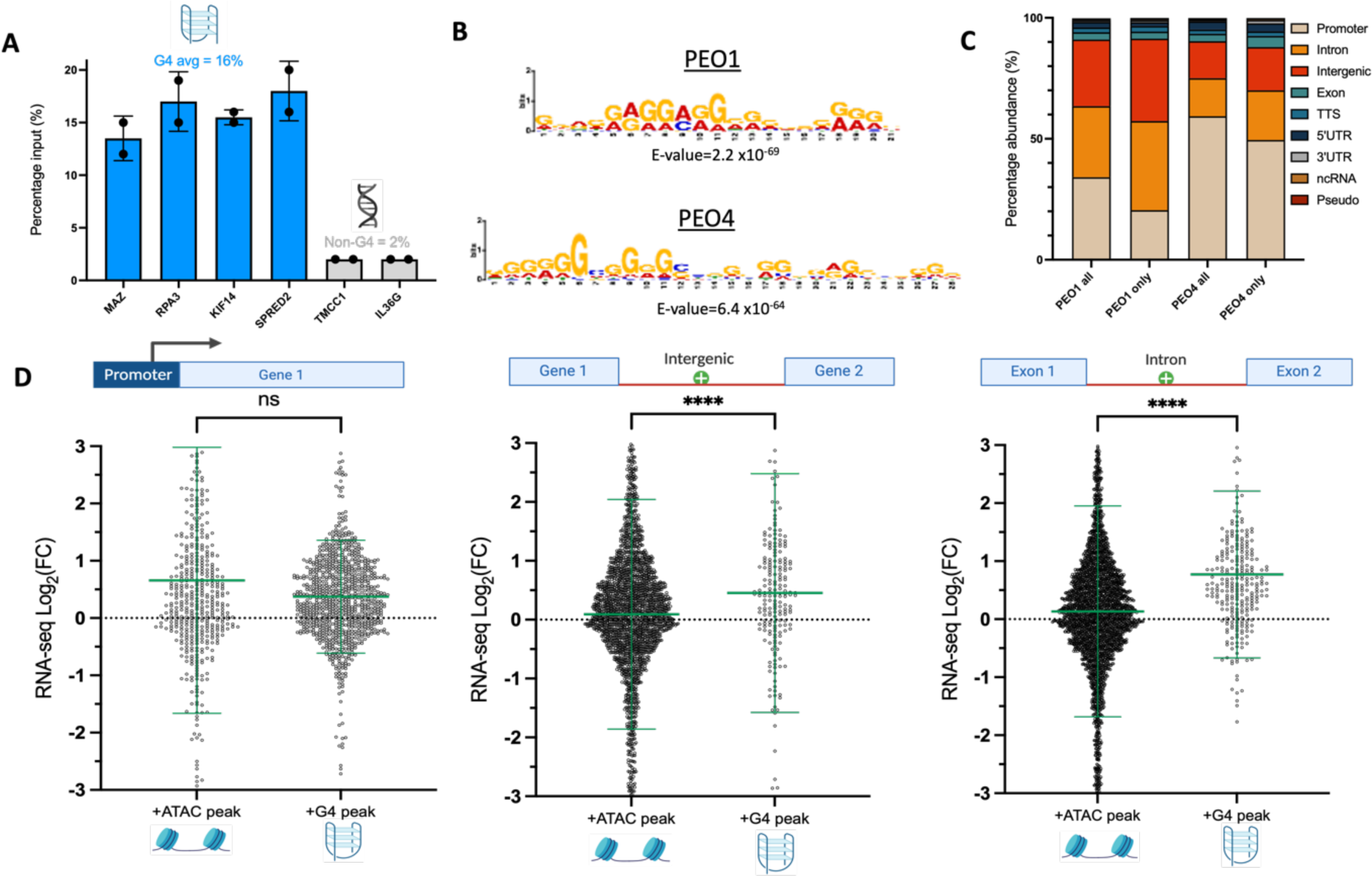
A) BG4 ChIP-qPCR with primers targeting G4 regions (MAZ, RPA3, KIF14, SPRED2) or non-G4 sites (TMCC1, IL36G) in chromatin extracted from PEO1 cells. BG4 immunoprecipitation results in an 8-fold enrichment for G4 regions over non-G4 sites. B) Most enriched sequence motif amongst top 1,000 G4 peaks in PEO1 and PEO4. Motif enrichment performed with MEME suite. C) Genomic distribution of BG4 ChIP-seq peaks in PEO1 and PEO4, annotated with HOMER. All peak locations were compared to peaks unique to only PEO1 or PEO4. D) Integration of ATAC, BG4-ChIP and RNA-seq data. Fold-change in expression (PEO4 relative to PEO1) of genes associated with gained ATAC peaks (that overlap putative G4 sequences) or gained G4 peaks identified with BG4 ChIP-seq. Comparisons are made for ATAC and G4 peaks located at promoters, introns and intergenic regions. Statistical significance assessed by Mann-Whitney U-test. Error bars show standard deviation of gene expression data collected in triplicate. FC=fold-change; ns = non-significant; **** = p<0.0001.

Interestingly, the total number of G4 peaks obtained in PEO4 cells was 2-fold lower when compared to PEO1 cells, with the latter producing 12,984 G4 peaks compared to 6,392 peaks in PEO4 cells. In agreement with previous G4 sequencing studies,^11,17,18,20^ the most enriched site of G4 formation was gene promoters– with 34% of PEO1 and 59% of PEO4 BG4 ChIP-Seq peaks residing within promoter regions (Figure 1C). However, when considering the distribution of G4 peaks that are unique to either PEO1 or PEO4, localisation at promoters dropped by approximately 10%, with a corresponding increase in abundance of G4s at intronic and intergenic regions.

We next asked if changes in G4 distribution between PEO1 and PEO4 cells were associated with differences in gene expression. To address this, we performed RNA-seq and linked the location of G4 peaks to the expression of the gene within the nearest transcription start site (termed G4 proximal or associated genes). Since G4s are known to form almost exclusively within open chromatin regions,^11^ we reasoned that any positive correlation between G4 formation and gene expression may be due to their enrichment at chromatin sites that are accessible for RNA polymerase binding. To account for this, we performed ATAC-seq (assay for transposase accessible chromatin) in PEO1 and PEO4, as previously described.^38^ Chromatin accessibility maps were then intersected with RNA-seq data, to directly compare changes in gene expression associated with increased chromatin accessibility versus G4 formation. To further control for the sequence bias associated with GC-rich G4 sites, we specifically selected ATAC peaks that overlapped with putative G4-forming sequences, obtained from *in vitro* G4-seq experiments (under K^+^ stabilisation).^39^ The expression of genes associated with these GC-rich ATAC-peaks was then compared to genes proximal to G4 regions immunoprecipitated by BG4 in either PEO1 or PEO4 cells.

Given the wealth of literature indicating G4 formation at promoter sites is associated with elevated gene expression,^11,17,18,20^ we initially limited our analysis to promoter G4s that appeared in PEO4 cells but were not detected in PEO1 cells. We found that genes associated with new promoter G4 peaks in PEO4 on average increased in expression (average Log_2_FC (fold-change) = +0.37, Figure 1D). However, this change was not significantly different from the increase in gene expression observed for promoter sites that acquired chromatin accessible regions detected by ATAC-seq (average Log_2_FC = +0.66, p=0.26, Figure 1D). This result contrasts with trends observed in previous genomic studies focused on different cell lines (e.g. healthy *vs* immortalised cells)^11^ and indicates that G4 formation at promoter regions of drug-resistant PEO4 cells does not confer a greater epigenetic advantage when compared to the simple gain of chromatin accessibility.

Nevertheless, a substantial body of work has indicated that non-promoter sites such as intergenic regions are of high relevance for the epigenetic regulation of drug-resistant ovarian cancer cells.^24–26^ Therefore, we next investigated whether the gain of G4 structures at non-promoter regions was associated with increased transcriptional levels when compared to the gain of accessible chromatin at the same loci. Specifically, we examined non-promoter regions that exhibited the highest G4 abundance in PEO1 and PEO4: intergenic regions and introns, which have previously been studied for their roles in epigenetic regulation.^40,41^ Strikingly, for both introns and intergenic regions, the formation of G4 structures was associated with a highly statistically significant increase in gene expression when compared to the gain of chromatin accessible sites in the same regions (p<0.0001, Figure 1D). More specifically, genes associated with new intergenic G4s had an average Log_2_FC 5-fold higher than genes associated with new chromatin accessible sites. Similarly, the gain of intronic G4s resulted in an average 6-fold higher increase in expression than the gain of ATAC peaks at introns (p<0.0001, Figure 1D).

These results indicate that non-promoter G4s, which have largely been overlooked in previous studies, may be of particular relevance to the epigenetic reprograming that enables ovarian cancer cells to become drug resistant. Furthermore, we observed that the loss of intergenic or intronic G4s was not associated with a statistically significant reduction in gene expression relative to the loss of chromatin accessible sites (Figure S2). This further suggests that the formation of G4 structures at introns and intergenic regions is transcriptionally functional and may be leveraged by cells to achieve higher levels of gene expression.

### The transcriptional associations of G4s are independent of BRCA2 status

Previously, it has been proposed that the difference in chemotherapy sensitivity between PEO1 and PEO4 was solely due to the different BRCA proficiency of the cells. PEO1 cells harbour a nonsense mutation in exon 11 of *BRCA2* leading to the loss of protein activity whereas PEO4 cells have a reversion mutation in *BRCA2* that restores the open reading frame.^42,43^ To disentangle the impact of BRCA status from the epigenetic rewiring that may contribute to drug-resistance, we additionally performed G4-mapping on PEO1 cells that acquired a spontaneous reversion mutation that restores BRCA2 function. These PEO1^BRCA+^ cells are thus expected to have similar capacity for homologous recombination to that of PEO4,^42^ and in turn are a useful model for independently assessing the epigenetic effects that may drive drug resistance.

Firstly, the BRCA2 status of both cell lines was confirmed with the RNA-seq dataset showing the key mutations in exon 11 of BRCA2 previously reported (Figure S3).^42^ MTS cell viability assays were used to compare the sensitivity of this pair of cell lines to cisplatin, revealing PEO1^BRCA+^ cells were approximately 6-fold more sensitive to cisplatin than PEO4 (Figure 2A). This is in agreement with previous literature reporting a 3-9 fold difference in sensitivity between PEO1^BRCA+^ and PEO4.^35,45^ This result thus suggests that BRCA2 status alone is not a reliable indicator of chemotherapy sensitivity and that transcriptional mis-regulation may instead be a key component in the development of drug resistance in ovarian cancer, as previously suggested.^4,24–26^

**Figure 2.**
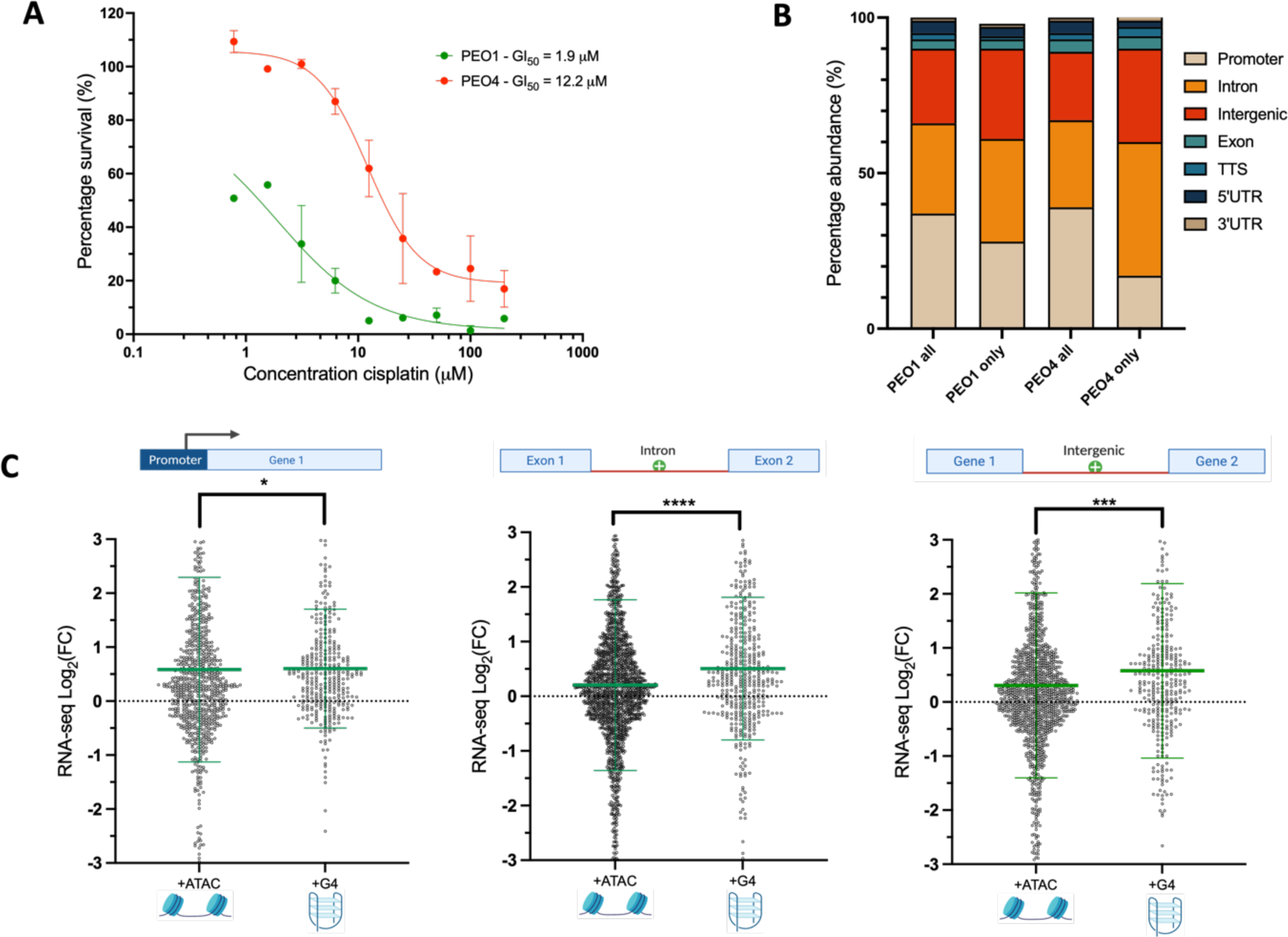
A) Cell viability of PEO1^BRCA+^ and PEO4 in increasing concentrations of cisplatin, treated across 72 hours. Error bars are standard deviation of results collected in triplicate. B) Genomic distribution of BG4 CUT&Tag peaks in PEO1^BRCA+^ and PEO4, annotated with HOMER. All G4-peaks detected in both PEO1/4 (PEO1/4-all) were compared to cell-type specific G4s (PEO1 only or PEO4 only). C) Integration of ATAC, G4 CUT&Tag and RNA-seq data in PEO1^BRCA+^/PEO4. Fold-change in expression (PEO4 relative to PEO1) of genes associated with gained ATAC peaks (that overlap putative G4 sequences) or gained G4 peaks identified with BG4 CUT&Tag. Comparisons are made for ATAC and G4 peaks located at promoters, introns and intergenic regions. Statistical significance assessed by Mann-Whitney U-test. Error bars show standard deviation of gene expression data collected in triplicate. FC= fold-change; *= p<0.05; *** = p<0.001 **** = p<0.0001.

We then sought to generate G4 maps in the PEO1^BRCA+^/PEO4 pair to further assess the relevance of G4s in the differential sensitivity against cisplatin observed in this cell line pair. Moreover, to ensure that the G4 maps generated were not biased by a specific G4-mapping technique used, we have detected G4s in this cell line pair using BG4 CUT&Tag rather than BG4 ChIP-Seq for comparison. Interestingly, the PEO1^BRCA+^/PEO4 pair also showed an overall reduction in G4 peaks in PEO4 cells (13,747 peaks in PEO1^BRCA+^ *vs* 8,178 in PEO4), with the overall most abundant detection of G4s in gene promoters. However, intergenic and intronic regions were still the most common site for detection of G4 peaks that were unique to PEO4 cells (Figure 2B). Thus, both the number and the genomic distribution of G4s detected by BG4 CUT&Tag in the PEO1^BRCA+^/PEO4 pair are similar to those found in the comparative analysis between PEO4 and parental PEO1 (i.e. *BRCA2* mutant) cells. These results further validate the consistency of the G4 maps obtained and highlights a potential regulatory role that these structures might play in establishing epigenetically driven resistance.

After integrating the ATAC-seq and RNA-seq data with the G4s maps obtained in the PEO1^BRCA+^/PEO4 pair, we observed a similar trend when considering the effects of promoter versus intergenic and intronic G4s on gene expression (Figure 2C). The appearance of new, PEO4-specific G4s at promoters resulted in a limited gain of gene expression compared to gain in chromatin accessibility (p=0.03, average Log_2_FC ATAC = +0.58, average Log_2_FC G4 = +0.60, Figure 2C). This is in contrast to the significant increase in expression of genes linked to the formation of new G4s at intergenic and intronic regions in PEO4 cells (p<0.001, Figure 2C). This data indicates that the global reduction of G4s in drug-resistant ovarian cancer cells co-occurs with a selective enrichment of G4s at intergenic regions and introns, which is in turn associated with elevated gene expression at these sites. Notably, these observations are independent of both the BRCA2 status of the ovarian cancer cells (BRCA wildtype or BRCA mutant) and the G4 mapping technique used (BG4 ChIP or G4-CUT&Tag). Altogether, these results strongly indicate a potential role of non-promoter G4s in the epigenetic regulation of genes altered after drug-treatment in ovarian cancers.

### Genes linked to G4 sites are important for drug responsiveness

We next investigated the biological pathways that were enriched in genes associated with new G4 peaks in PEO4 and performed KEGG pathway enrichment in the PEO1^BRCA+^/PEO4 cells using the ShinyGO gene-set enrichment tool.^46^ Pathway enrichment of upregulated genes in proximity to PEO4-specific intronic and intergenic G4s highlighted multiple pathways (Figure S4). Of particular interest were the WNT, hippo and calcium signalling pathways, all of which have been strongly linked to the epithelial to mesenchymal transition (EMT).^47–49^ During EMT, cells are commonly epigenetically reprogrammed to a progenitor-like state which is accompanied by increased proliferation and reduced rates of apoptosis.^49–51^ The importance of EMT in acquired drug resistance has been well documented in many cancers and has been shown to be of high importance in the context of drug-resistant ovarian cancer.^50,51^

Interestingly, there was also a particularly high enrichment of genes at the beginning of the WNT signalling pathway (Figure S5), where perturbations may have larger downstream effects. This finding was further validated by gene set enrichment analysis (GSEA),^52^ which revealed that EMT, as well as the WNT-beta catenin pathways were amongst the significantly enriched hallmarks for upregulated genes associated with PEO4-specific intergenic and intronic G4s (Table S1). From this result, we hypothesised that G4 formation at key non-promoter sites may be leveraged by cells to maintain a high expression level of such resistance-relevant signalling pathways.

To further investigate whether genes associated with non-promoter G4s have previously been linked to drug-resistance, we performed a literature search with NDEx iquery, a neural network-based tool that searches for geneset similarities across >80,000 publications.^53^ The publication that had the highest similarity to upregulated genes associated with PEO4-specific intergenic and intronic G4s was highly relevant to drug-resistant ovarian cancer, and specifically focused on how increased transcription correlates with loss of DNA methylation in cisplatin-resistant ovarian cancer cells (Table S2).^54^ Importantly, this study was based on an alternative drug-sensitive and drug-resistant ovarian cancer cell line pair named M019 and M019i.^54^ Similarly to our analysis, the authors found that the primary methylation changes that predicted gene expression occurred at intergenic and intronic regions. Given recent reports that G4 formation actively inhibits DNA methyltransferase activity,^55,56^ this overlap may suggest a mechanism by which G4 formation could be leveraged by ovarian cancer cells to achieve epigenetic resistance to chemotherapy. It was also found that WNT signaling was amongst the most significantly altered gene pathways between the M019 and M019i pair,^54^ which further exemplifies the significance of this pathway in the development of drug-resistant ovarian cancer independently of the cell model used.

In contrast to genes associated with intergenic and intronic G4s, genes that acquired new promoter G4s in PEO4 were not significantly enriched for any KEGG pathways, which again highlights the nominal relevance of G4 formation at promoter sites in this drug-resistant cell model. Similarly, when utilising NDEx iquery to identify previously reported genesets that significantly overlapped with promoter G4-linked genes, none of the highlighted publications had relevance to either ovarian cancer or drug sensitivity (Table S3). The pathway enrichment results thus provide compelling evidence that the impact of G4 formation in chemo-resistant ovarian cancer cells is location-specific and highly focused around non-promoter intergenic and intronic sites, that are associated with the expression of key drivers of drug resistance.

### G4s act cooperatively with transcriptional enhancers and one-another

The epigenetic state of a cell is coordinated by multiple genomic sites that recruit key transcription factors, coactivators and chromatin remodelling complexes.^57,58^ It is therefore conceivable that the formation of G4s may synergise with or be linked to other epigenetic marks. In particular, transcriptional enhancers have been noted in recent years to be critical members of the epigenetic toolkit of a cell – allowing cells to rapidly control the expression of genes across thousands of base pairs.^40,59^ Importantly, epigenetic regulation mediated by transcriptional enhancers has been shown to be of particular relevance in ovarian cancer and drug resistance development.^25,26,60^ To assess if cooperation between G4s and enhancers could be relevant in these ovarian cancer cells, we performed CUT&Tag in PEO1 and PEO4 cells on the histone mark H3K27ac, commonly used to assign transcriptional enhancer sites.^61^ Specifically, we characterised the PEO1^BRCA+^/PEO4 pair to remove confounding, non-epigenetic effects that may be attributed to differences in BRCA2 status.

Genome-wide mapping of the histone mark H3K27ac revealed that the majority of enhancers are detected in non-promoter regions, with 50% of PEO4-specific enhancers being found at introns and 33% at intergenic regions compared to only 7% detected in promoters (Figure 3A). This again highlights the importance of non-promoter regions for defining the epigenetic landscape of drug-resistant ovarian cancer cells. Strikingly, there was a large overlap of enhancers and G4s, with 84% of G4s in PEO1 and 78% of G4s in PEO4 overlapping with transcriptional enhancers. Despite this, there were also many enhancer sites where G4s were absent, given that there were substantially more enhancers detected than G4s (24,647 enhancers in PEO1^BRCA+^ and 27,647 in PEO4). We thus tested if the presence of G4s within transcriptional enhancer regions resulted in elevated gene expression when compared to an average enhancer. To control for GC-richness, enhancers that overlapped with putative G4-forming sequences (OQS, defined with *in vitro* G4-seq)^39^ were compared to enhancers that coincided with validated G4 peaks detected by BG4 CUT&Tag. This investigation revealed that genes proximal to enhancers that contained G4 peaks displayed a significantly higher level of gene expression when compared to genes proximal to an average, GC-rich enhancer (average enhancer Log_2_FC = +0.73, average G4 enhancer Log_2_FC= +1.00, p<0.0001, Figure 3B), indicating that G4-containing enhancers represent a subcategory of particularly potent enhancer sites.

**Figure 3.**
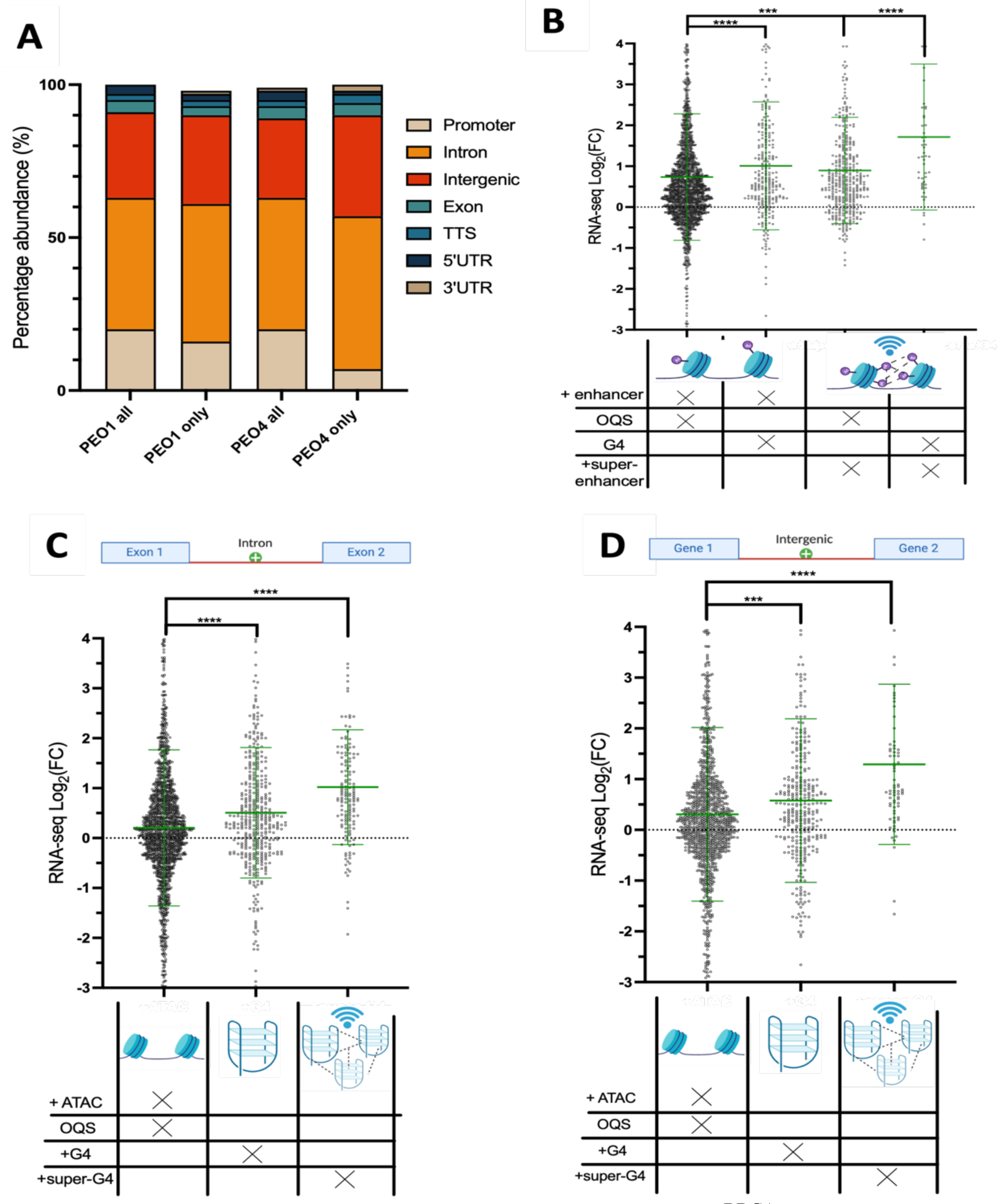
A) Genomic locations of enhancer peaks in PEO1^BRCA+^ and PEO4, identified by H3K27ac CUT&Tag. The distribution of all peaks in either cell line is compared to peaks specific to either PEO1 or PEO4. B) Fold-change in gene expression (PEO4 relative to PEO1) of genes associated with gained enhancers and super-enhancers. Enhancers are defined by H3K27ac peaks and super-enhancers by significant clusters of enhancers found in a 12.5 kb window, identified with the ranked ordering of super-enhancers (ROSE) algorithm. ^62,63^ Genes associated with gained enhancer/super-enhancer peaks that overlap a putative G4 sequence (OQS) are compared to genes proximal to enhancers that overlap G4 peaks identified with BG4 CUT&Tag. C) Change of expression for genes associated with super-G4s regions, defined by ROSE as significant clusters of G4 CUT&Tag peaks at intronic or D) intergenic regions. The expression of super-G4 linked genes is compared to genes associated with singular G4s or ATAC-peaks that overlap OQS. Statistical significance assessed by Mann-Whitney U-test. Error bars are standard deviation of gene expression measured across 3 biological replicates. FC= fold-change; ns = non-significant; *** = p<0.001, **** = p<0.0001.

Enhancers are additionally known to act cooperatively with one another where clusters of enhancer regions create “super-enhancer” sites with increased regulatory power.^64,65^ Super-enhancers have been well established to be some of the most efficient epigenetic regulators within cells, being particularly important for controlling the expression of master transcription factors and driving cell fate.^63–65^ Moreover, increasing evidence has established the role of super enhancers in enabling cancer cells to rapidly change expression profiles in response to drug treatment.^24–26^ The superior effects of super-enhancers are thought to be achieved partly by their ability to recruit transcription-regulating protein with intrinsically disordered domains prone to liquid-liquid phase separation.^33,34^ Similarly, G4s are validated binding partners of multiple transcription factors and co-activator proteins^19^ and can themselves trigger phase-separation events by nucleating intermolecular strand interactions.^32,66,67^ We therefore questioned whether clusters of G4s may act similarly to super-enhancers.

To first identify super-enhancers, we used the ranked ordering of super-enhancers (ROSE) algorithm to identify clusters of H3K27ac peaks within 12.5 kb of one another, as previously described.^62,63^ We found 1,235 super-enhancer sites that were specific to PEO4 cells and not found in PEO1. As expected, genes associated with PEO4 super-enhancer sites displayed a statistically significant increase in gene expression when compared to genes proximal to individual enhancers (super-enhancer genes: average Log_2_FC= +0.89, enhancer genes: average Log_2_FC= +0.73, p<0.001, Figure 3B). The expression levels of genes regulated by PEO4-specific super-enhancers were additionally increased when a given super-enhancer overlapped with a G4 peak, again indicating regulatory cooperation between enhancers and G4s.

We next applied the ROSE algorithm to the G4 CUT&Tag data, to identify clusters of G4 peaks that we coin “super-G4” sites, which have not previously been investigated. We found that 636 new super-G4s were acquired in PEO4 cells, with an average of 4.2 G4 peaks spanning 16 kb Surprisingly, we observed that genes proximal to intronic and intergenic super-G4s displayed increases in gene expression even greater than that observed for genes linked to super-enhancers. Genes linked to intergenic and intronic super-G4 sites had an average Log_2_FC= +1.29 and +1.02 respectively, compared to super-enhancer associated genes with Log_2_FC=+0.89 (Figures 3B-D). This demonstrates that whilst individual enhancers have larger effects on gene expression than individual G4 sites, the cumulative effects of multiple closely situated G4s is greater than that of super-enhancers. Additionally, the majority of genes (65%) associated with super-G4s were not associated with super-enhancers, indicating super-G4s may regulate a unique set of gene pathways, thus making the potential targeting of G4 for epigenetically re-wiring of drug resistant cancer cells orthogonal to current epigenetic strategies deployed in the clinic. Given the undoubtable relevance of super-enhancers in defining transcriptional states within cells, these results indicate that super-G4s may have a previously unconsidered role in regulating gene expression in drug-resistant cells, which rivals or even surpasses the level of transcriptional enhancement achieved by super-enhancers.

### The epigenetic effects of non-promoter G4s are observed in additional patient-derived ovarian cancer cell lines

Multiple pairs of drug-sensitive and drug-resistant ovarian cancer cell lines have been previously established and investigated to understand epigenetic mechanisms that drive acquired drug tolerance.^68–70^ We thus expanded our analysis of G4-mediated regulation of drug resistance to other ovarian cancer models, with the aim of assessing whether the correlations of gene expression and G4 formation was limited to the PEO1/PEO4 cell line pair. To achieve this, we reasoned that the gain of accessible sites with a putative G4 forming sequence may be used to predict increases in G4 formation in drug-resistant cells. Whilst not all G4 sequences in accessible sites necessarily fold into G4s within cells,^11^ chromatin accessibility is a pre-requisite for G4 formation and thus the two states are intimately linked.^11^ We therefore intersected sites with altered chromatin accessibility (identified by ATAC-seq) with putative G4-forming sequences identified by *in vitro* G4-seq.^39^

To test the validity of this approach, we first intersected ATAC-seq, G4-seq and RNA-seq data in PEO1 and PEO4, for which we had attained G4 maps with two different genomic methods and as described above. More specifically, we compared the expression of genes that had gained a chromatin accessible site in resistant cells, with those that gained a chromatin-accessible site that also coincided with a putative G4 forming sequence characterised by G4-seq. From this analysis we found the presence of G4 sequences at promoters did not cause a significant change of associated gene expression (p=0.57, Figure 4A). In contrast, G4-forming sequences at accessible intergenic and intronic sites resulted in a significant increase in expression of nearby genes (p=0.0002 and p=0.0001 respectively, Figure 4A). This result demonstrates that the combination of ATAC, G4-seq and RNA-seq is sufficient to predict G4-mediated effects at specific genomic loci, yielding results that are highly comparable to trends found with genomic G4 maps obtained with either BG4 ChIP-seq or CUT&tag.

**Figure 4.**
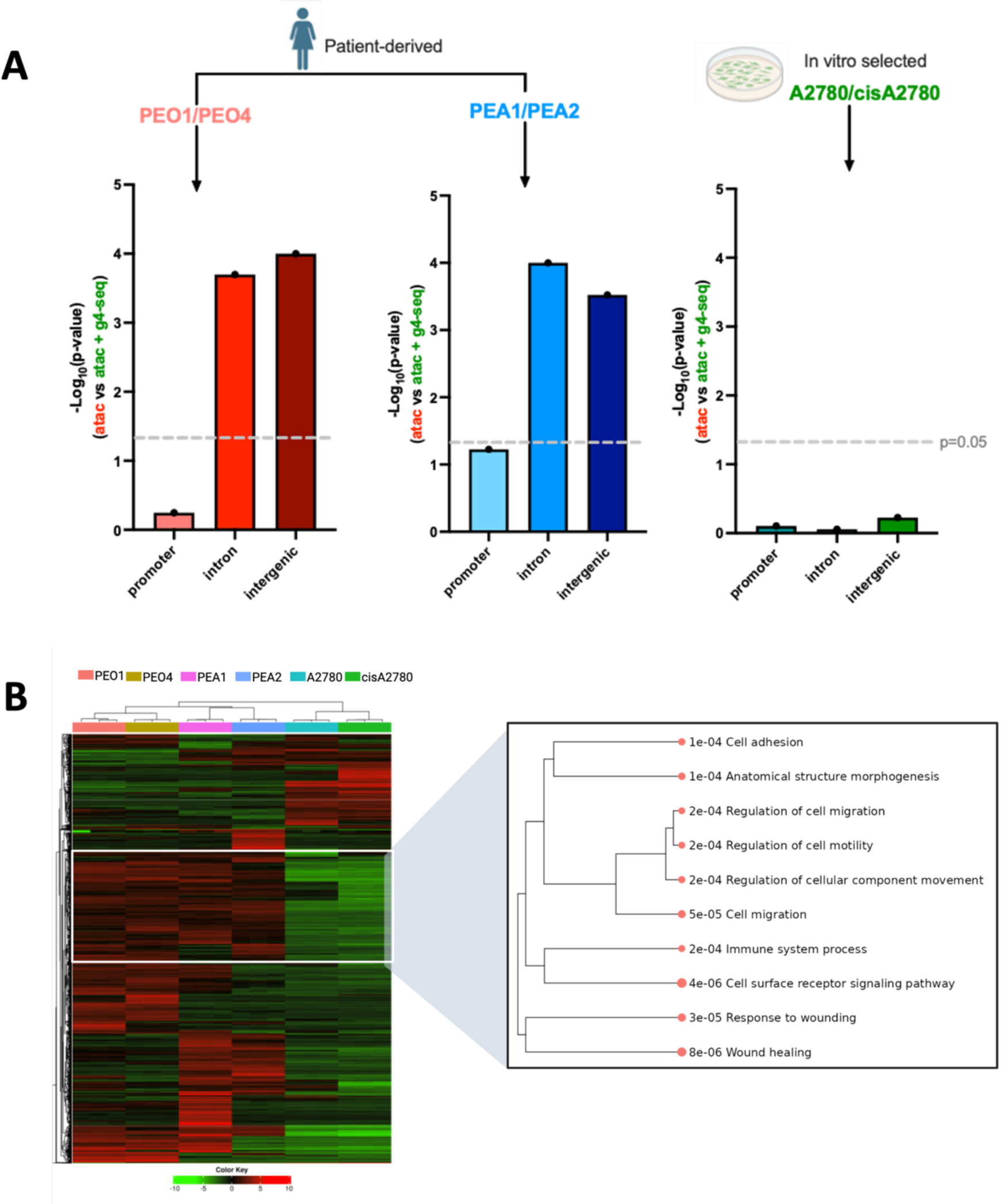
A) Comparison of the effects of putative G4 sequences on gene expression within 3 paired drug-sensitive and drug-resistant ovarian cancer cell lines: PEO1 and PEO4, PEA1 and PEA2, A2780 and cisA2780. The expression of genes associated with gained ATAC peaks was compared to genes that gained ATAC peaks also overlapping putative G4-sequences (detected by G4-seq) at promoters, introns and intergenic regions. Differences in expression between the two groups was assessed by p-value obtained from Mann-Whitney U-test. Dotted line at p=0.05. B) Gene expression heat map of top 1,000 most differentially expressed genes between PEO1, PEO4, PEA1, PEA2, A2780 and cisA2780, based on previously acquired RNA-seq data.^24^ Samples clustered by average Pearson correlation. Pathway enrichment was conducted on genes uniquely expressed in PEO1/PEO4 and PEA1/PEA2 compared to A2780/cisA2780 revealing genes enriched in cell adhesion, motility and wound healing. Analysis performed in iDEP.^71^

Encouraged by these observations, we next decided to extend our analysis to other pairs of drug-sensitive and drug-resistant cell lines that have available ATAC-seq and RNA-seq data. We included the patient-derived cell lines PEA1 and PEA2, which similar to the PEO pair were derived from a single high-grade serous ovarian cancer (HGSOC) patient before and after she developed resistance to platinum-based chemotherapy (cisplatin).^69^ However, unlike PEO1, PEA1 cells are treatment naïve and were established before the patient received any chemotherapy. We additionally investigated the A2780 ovarian cancer cell line and its cisplatin-resistant pair cisA2780.^68^ In contrast to the PEO and PEA pair, the cisA2780 cell line was established by exposure of the parent cell line to extended, high doses of cisplatin *in vitro*.^68^ Therefore, it has been suggested that the mechanisms by which these cells have developed drug resistance may be distinct to those that occur natively *in vivo*,^24^ where factors such as the tumour microenvironment and pharmacokinetics may play a more prominent role.

Performing the same integration analysis of ATAC-seq, G4-seq and RNA-seq data in PEA1/PEA2 revealed that genes nearby newly accessible sites that contained putative G4-forming sequences had greater increases in expression, with this difference being statistically significant at intergenic and intronic regions (p<0.0001 and p=0.003 respectively, Figure 4A). However, the effects of G4 sequences at promoters only had borderline significance (p=0.06, Figure 4A), similarly to that observed for PEO1 and PEO4 cells. Furthermore, genes that gained putative intergenic/intronic G4 sites in PEA2 cells were enriched in EMT signatures (GSEA: FDR = 2.7e^-10^), in addition to KEGG pathways such as MAPK signalling and transcriptional mis-regulation (Table S4, Figure S6). However, no pathways were significantly enriched for genes linked to new putative promoter G4 sites in PEA2 cells. This result therefore corroborates the finding that non-promoter G4s contribute to regulation of resistance-related pathways in drug-resistant ovarian cancer cells obtained from different patients, suggesting that G4 formation at key regulatory sites could be a common epigenetic mechanism to acquire drug resistance in ovarian cancer patients.

In contrast, when the same analysis was extended to the A2780/cisA2780 pair, no significant effect of G4 sequences was observed at any of the tested genomic sites (Figure 4A). To further probe differences between the patient-derived and *in vitro* selected drug-resistant cells, the RNA-seq data was further investigated. Whilst each cell line pair had distinct expression profiles, sample clustering and principal component analysis revealed the PEO and PEA pairs clustered together relative to the A2780 pair (Figure 4B, S7). This result also aligns with previous work showing that changes in chromatin accessibility are more comparable between patient-derived resistant cells than those established artificially *in vitro*.^24^ Clusters of genes that showed high expression in PEO/PEA cells, but not A2780 cells were found to be enriched in processes such as cell adhesion, migration, immune system responses and wound healing (Figure 4B). These pathways are less likely to be activated *in vitro* where cells are removed from the tumour microenvironment and indicate that the cisA2780 cell line likely has unique adaption pathways that are seemingly not dependent on G4 formation. Moreover, the A2780 cell line has previously been shown to harbour a distinct genetic profile compared to patient-derived HGSOC samples, which may make it a poor model for epigenetic changes relevant to ovarian cancer patients.^72,73^

Together, these results indicate that G4s may play a fundamental role in the distal regulation of resistance-related genes that is not limited to the context of a single patient. Additionally, it highlights the important differences between cell models that are established *in vivo* and *in vitro* – the latter of which may not effectively reconstitute the epigenetic changes that occurs naturally in patients.

### Small-molecule targeting of G4s re-sensitises drug-resistant cells via epigenetic reprogramming

Given the strong, location-specific correlations we found between G4s and gene expression within drug-resistant ovarian cancer cells, we hypothesised that direct targeting of G4s may be used to disrupt the epigenetic networks that drive drug resistance. To investigate this, we first examined if treating cells with the G4-stabilising small molecule pyridostatin (PDS),^74^ had any effect on the cisplatin sensitivity of PEO1^BRCA+^ or PEO4. This experiment was conducted using a single dose of PDS (0.5 μM) that did not induce any toxicity in either PEO1 or PEO4 after 72-hour incubation alone (Figure S8). The PDS-treated cells were simultaneously exposed to increasing concentrations of cisplatin to test if the toxicity of the chemotherapy agent was increased in the presence of the G4 binder. Interestingly, we found that co-treating cells with PDS and cisplatin had no significant effects on the viability of PEO1 cells but resulted in a large 3-fold increase in cisplatin sensitivity for PEO4 cells (Figure 5A), suggesting that PDS can selectively increase cisplatin toxicity in drug-resistant PEO4 cells.

**Figure 5.**
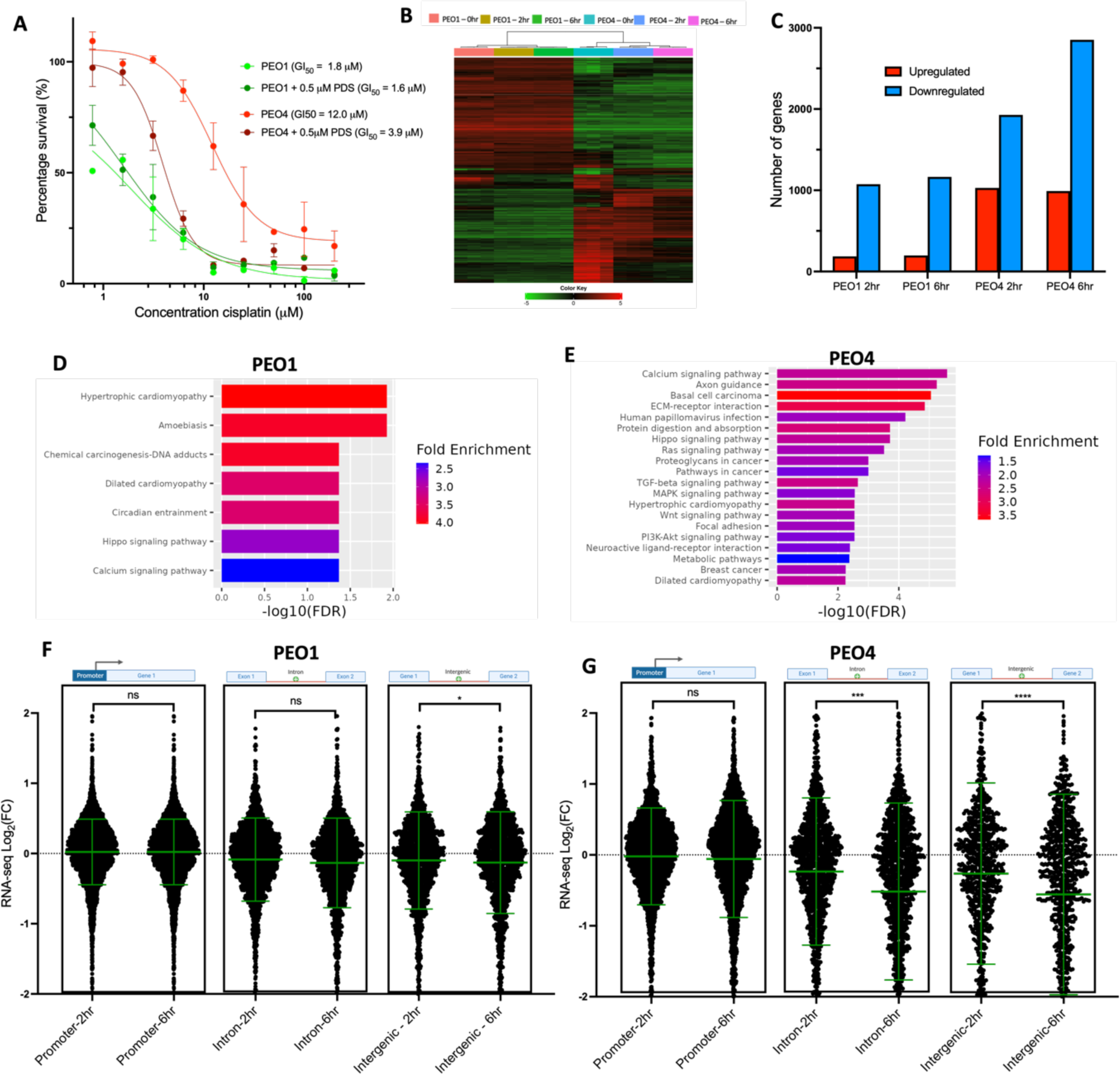
A) Cell viability of PEO1^BRCA+^ and PEO4 in response to increasing concentrations of cisplatin with or without cotreatment with G4 ligand PDS (0.5 μM), incubated for 72 hours. PDS significantly re-sensitises PEO4 cells to cisplatin but has no effect on PEO1. Error bars are standard deviation of results collected in triplicate. B) Heat map of top 1,000 most differentially expressed genes in PEO1 and PEO4 after, 0, 2 and 6 hours of PDS treatment (5 μM). C) Number of significantly differentially expressed genes (|FC|>1, FDR<0.05) in PEO1 and PEO4 after 2 and 6 hours of PDS (5 μM) treatment. D) KEGG pathway enrichment of genes significantly downregulated in PEO1^BRCA+^ and E) PEO4 after 6 hours PDS (5 μM) treatment. F) Fold-change in gene expression for genes associated with G4 peaks (identified by BG4 CUT&Tag), after treatment with PDS (5 μM) for 2 or 6 hours. Expression is compared for genes proximal to G4 peaks at promoters, introns and intergenic regions in PEO1^BRCA+^ and G) PEO4. Statistical significance assessed by Mann-Whitney U-test. Error bars are standard deviation of gene expression data acquired in triplicate. FC= fold-change; ns = non-significant; *= p<0.05, *** = p<0.001, **** = p<0.0001.

Previously, PDS has been reported to cause DNA damage at G4 sites in proportion to the number of accessible G4 regions and thus it is expected to synergise with DNA cross-linking drugs such as cisplatin.^75,76^ However, given no significant effect was observed within PEO1 cells that contain ∼2-fold more G4s than PEO4, the DNA-damaging effects of PDS do not explain the selective, partial re-sensitisation of PEO4 cells. Instead, we reasoned that PEO4 cells have a unique epigenetic state that is more reliant on the presence of G4s and super-G4s at specific genomic locations, such as introns and intergenic regions. To test this hypothesis, we used RNA-seq to map the transcriptional profiles of PEO1 ^BRCA+^ and PEO4 cells after treatment with PDS for 2 or 6 hours at approximately the 72-hour IC_50_ concentration (5 μM, Figure S8).

RNA-seq data revealed that the transcriptional profiles of the cells were substantially altered by PDS treatment, particularly in PEO4 (Figures 5B, C). For both PEO1 ^BRCA+^ and PEO4, the majority of differentially expressed genes (|FC|>1, FDR<0.05) were down-regulated (Figure 5C). For instance, after 6 hours of PDS treatment 2,851 genes in PEO4 were significantly downregulated compared to 990 upregulated genes. In contrast 1,163 genes were downregulated and 198 genes upregulated in PEO1^BRCA+^ after 6 hours. This aligns with previous studies showing that G4 targeting with small molecules results in global downregulation of gene expression, which may be due to DNA damage caused by hyper-stabilising naturally dynamic G4 structures, or by displacing the endogenous binding partners of G4s that have been shown to promote gene expression.^21,77^ Additionally, the transcriptional effects PDS had in PEO1 cells were mostly plateaued within 2 hours, with the 6 hour treatment condition producing limited additional changes (15 genes significantly altered between 2 and 6 hours in PEO1). However, there were >1,000 genes significantly altered between the 2- and 6-hour treatment timepoint for PEO4. This demonstrates that in PEO4 the transcription-altering effects of PDS are not only greater in magnitude, but also have stronger temporal dependence, suggesting that perturbing G4-prevalence on PEO4 cells has a more global impact on gene expression changes that might reflect a more relevant role of G4s structures in the maintenance of transcriptional homeostasis in PEO4 cells.

To understand which gene pathways were significantly perturbed by G4 stabilisation we performed pathway enrichment analysis on significantly altered genes. Across all conditions, multiple KEGG pathways were significantly affected (Figures 5D, E). In PEO4, many signalling pathways were enriched amongst downregulated genes, including Hippo, WNT, MAPK and TGF-B (Figure 5E), all of which are implicated in the control of EMT, responses to stress and are reported to contribute to acquired drug tolerance in ovarian cancer.^50,78,79^ The enrichment of signalling pathways in altered PEO4 genes may also explain the temporal effect of G4 stabilisation, as these pathways typically regulate the localisation of master regulatory proteins which in turn affect the expression of multiple downstream genes over longer time periods.^80^ Additionally, multiple pathways contributing to cancer proliferation including those found in basal cell carcinomas and breast cancers were down-regulated, suggesting PDS treatment may result in a global reduction of cancer cell fitness. In comparison, significantly fewer signalling pathways were enriched for down-regulated genes in PEO1, with top hits being in pathways such as cardiomyopathy and processing of DNA adducts.

We next set out to link changes in gene expression post-PDS treatment to the presence of G4s at promoters, introns and intergenic regions as detected by BG4 CUT&Tag. By analysing the expression change of genes associated with promoter G4s, we observed that for both time points and cell lines, the average change in expression was negligible (Figure 5F,G). In PEO1^BRCA+^, G4 promoter-linked genes had an average Log_2_FC=+0.02 after 6 hours PDS treatment and in PEO4 an average Log_2_FC=-0.06. In contrast, genes linked to intergenic and intronic G4s had much larger changes of gene expression, particularly in the case of PEO4 where the effect was also time-dependent (Figures 5F,G). Specifically, after 6 hours PDS treatment, genes in PEO4 associated with intergenic and intronic G4s had an average Log_2_FC=-0.56. and -0.52 respectively, almost 10-fold greater than that of promoter G4-associated genes. We next examined the pathways associated with down-regulated genes linked to PEO4 intergenic/intronic G4s, and again found enrichment of WNT, Hippo and calcium signalling which indicates that these key pathways are epigenetically perturbed by G4 stabilisation (Figure S9). In contrast, no KEGG pathways were significantly enriched for down-regulated genes associated with promoter G4s, which further highlights that promoter G4s are unlikely to regulate any critical pathways associated with the development of drug resistance in ovarian cancer patients exposed to chemotherapy.

Altogether these experiments reinforce the notion that, in the context of PEO4 cells, the presence of non-promoter G4s are key for rewiring transcriptional profiles that might be linked to development of drug-resistance, whereas G4s forming at promoters have limited relevance. Furthermore, it supports our hypothesis that the specific re-sensitization of PEO4 cells with PDS is attributable to the stronger reliance that drug-resistant cells have on G4 structures to maintain a defined epigenetic state. This is evidenced by the greater global change of gene expression in PEO4 after G4 stabilisation, as well as the significantly higher association of these changes with the presence of G4s at intergenic and intronic sites. Overall, this highlights the potential of using G4 ligands to alter transcriptional pathways that are essential for ovarian cancer cells to acquire drug resistance and the potential for future exploration of G4 stabilisation to combat resistance acquisition.

## Discussion

DNA G-quadruplex structures have been widely linked to the epigenetic state of cells, with multiple independent studies showing endogenous G4 formation is correlated with overall increased gene expression.^11,17–20^ Given the enrichment of G4s within key oncogenes and globally in cancer cells,^10–14^ it has been speculated that G4 formation may contribute to transcriptional programs that promote proliferation during oncogenesis. However, cancer cells are not epigenetically stagnant but constantly evolving,^81,82^ leading to dynamic changes in gene expression alongside DNA structures such as G4s.^83^ In ovarian cancer and several other cancers, such epigenetic plasticity can lead to acquired drug resistance that is fatal to patients in most cases.^3,84,85^ Despite this, a systematic study to associate G4 formation and changes in transcription associated with drug resistant cells has not been previously performed, making it impossible to anticipate whether G4 formation may contribute to the remarkable and devastating capacity for drug-induced evolution that is typical of cancer cells.

In this work, we explored the impact that G4 formation has on the epigenetic state of drug-resistant ovarian cancer cells by generating G4 maps via two independent genomic methods (BG4 CUT&Tag and BG4 ChIP-Seq) in paired drug-sensitive and drug-resistant ovarian cancer cell lines (PEO1 and PEO4). We intersected G4 maps with maps of the transcriptional enhancer mark H3K27ac (CUT&Tag), chromatin accessible sites (ATAC-Seq) and gene expression changes (RNA-Seq) in the same cells to establish associations between G4 formation and global epigenetic changes linked to ovarian cancer drug resistance. Surprisingly, we found no effect of G4 formation when compared to increased chromatin accessibility at promoters, which is a stark difference from literature reports of the strong transcriptional effects of G4s forming within promoter regions.^11,17–20^

In contrast, we observed a highly significant association between G4 formation at introns and intergenic regions and increased gene expression. Generally, intergenic transcriptional enhancers have been linked to the regulation of house-keeping genes, whereas the targets of intronic enhancers are enriched for cell-type specific genes;^41,86,87^ however, both locations share the critical ability to regulate gene expression over long genomic distances. The location-specific association of G4s at introns and intergenic regions may therefore indicate that G4s act via distal regulatory mechanisms in drug-resistant cells, which may represent a distinct mechanism to the previously established impact of G4s at promoters.^11,17–20^ Crucially, this effect of G4s outside of promoters was also observed in a distinct pair of patient-derived cell lines, suggesting a common mechanism of epigenetic rewiring that leverages G4 formation at non-promoter sites to achieve transcriptional mis-regulation that leads to drug-resistance. Moreover, this association between G4s and gene expression was not detected in drug-resistant cells artificially induced *in vitro*, which are likely to be driven by distinct mechanisms which are not reliant on the tumour microenvironment.

We additionally found that the presence of G4s synergised with enhancers marked by H3K27ac, which were also predominantly found outside of promoters. Critically, we observed that significant clusters of G4 peaks, that we coin “super-G4s”, were associated with changes in gene expression even greater than that of super-enhancers, which are well established to be some of the most powerful regulatory elements in the genome.^63–65^ This result suggests that clusters of G4s may themselves act as distinct transcriptional super-enhancer regions, which has not been previously considered. We reason that the high binding affinity of G4s to several regulatory proteins,^19,28,30^ along with the ability of G4s to alter methylation profiles^55^ and to trigger phase separation events^32,66,67^ may make these structures particularly powerful effectors of long-range gene regulation outside of promoters (Figure 6).

**Figure 6.**
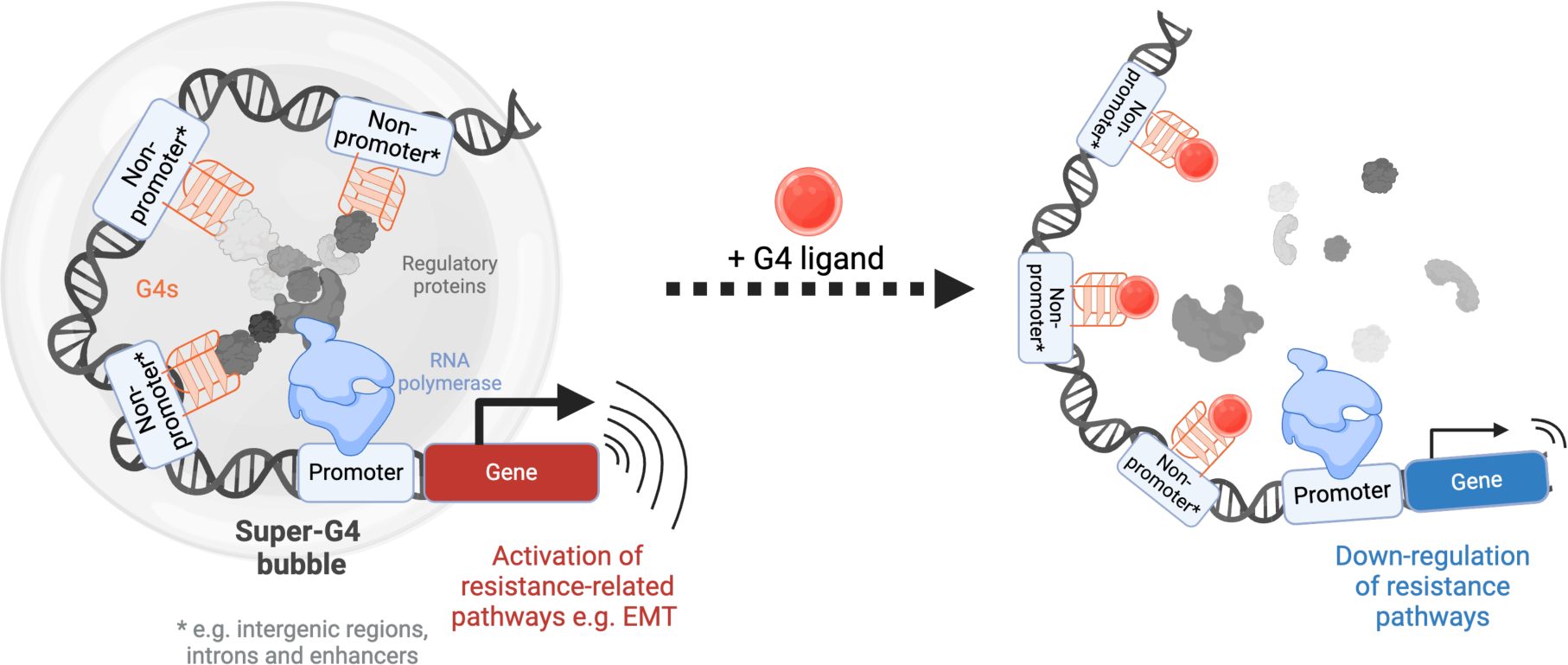
Proposed mechanism of distal, G4-mediated regulation of gene expression in drug-resistant ovarian cancer cells. Clusters of G4s (super-G4s) form at non-promoter sites and may recruit key regulatory proteins or trigger phase-separation events that increase expression of signalling proteins involved in drug responses. Incubation of cells with a G4 ligand (PDS) results in significant down-regulation of genes associated with drug resistance and leads to a marked increase in drug sensitivity.

The impact of such long-range regulation has begun to be unveiled by previous studies demonstrating that ovarian cancer cells can achieve larger transcriptional change by altering distal regulatory sites, rather than promoter regions.^24–26^ Perhaps this is because distal transcription-enhancing regions are able to regulate the expression of multiple genes at once, for instance, by creation of super-enhancer (or potentially super-G4s) bubbles which sequester multiple transcription-promoting proteins in a confined space.^33,34^ This may allow cells to rapidly change the expression of large groups of genes rather than on an individual basis and in turn lead to more efficient adaption to drug treatment.

To understand if disrupting long-range G4 networks may revert drug tolerant cells to a sensitive state, we investigated the effects of targeting G4s with the highly selective small-molecule PDS. Through G4 stabilisation we were able to induce significant changes in the transcriptional profile of drug-resistant cells, specifically for genes associated with intergenic and intronic G4s. Overall, global downregulation of genes was observed, particularly in signalling pathways such as WNT, hippo and calcium signalling proteins which have previously been implicated in EMT and acquired drug resistance of ovarian cancer cells.^47–51^ These transcriptional changes were accompanied by a significant increase in sensitivity of drug-resistant cells, but minimal changes in the response of drug-sensitive cells. The fact that G4 targeting specifically re-sensitises drug-resistant cells may allow for G4 ligands to be used to epigenetically reprogramme drug-resistant cells without altering the behaviour of drug-sensitive or potentially healthy cells (Figure 6). Currently, a small number of G4 ligands have entered clinical trials for cancer therapy^88–90^ and may therefore be ideal candidates for investigating the feasibility of G4 targeting to counter drug resistance. In the future, extending analysis of G4-mediated regulation beyond promoter sites may lead to further insights into the important role of these structures in the epigenetic evolution of cancer cells, potentially beyond ovarian cancer.

### Conclusions

In this work we have deployed a combination of genomics methods to underpin the link between G4 formation and epigenetic changes driving drug-resistant ovarian cancers. We have shown that G4s in promoter regions are surprisingly non-relevant to transcriptional changes key to epigenetically induced drug-resistance in patients, whilst G4s forming in intergenic and intronic regions provide a transcriptional advantage to such key resistance-associated genes. Additionally, our work demonstrated that small-molecule targeting of G4s in introns and intergenic regions can induce transcriptional changes that resensitise cells to chemotherapy agents, underscoring the potential of including G4 targeting in combination therapies for drug-resistant cancer treatment. Finally, we demonstrated that G4s and enhancers act synergistically, leading to increased transcriptional activity. This is further corroborated by the observation that cluster of G4s, which we have coined super-G4s, also act co-operatively and lead to synergistic enhancement of gene expression, similarly to the effects of super-enhancers. Importantly, genes and pathways that are regulated by super-G4s seems to be distinct to those regulated by super enhancers, indicating that super-G4 might represent an alternative epigenetic mark to enhance gene expression. Our work supports the relevance of G4 formation in epigenetic regulation and maintenance of chromatin architecture, stretching beyond promoters. Based on our study, we anticipate that the development of clinically viable G4 ligands could be a cornerstone for treatment of drug-resistant ovarian cancers.

## Methods

PEO1 and PEO4 cells (ECACC) were grown in RPMI 1640 media with Glutamax (Gibco) supplemented with 10% FBS.

### BG4 expression

*TES buffer*: 50 mM Tris-HCl, 1 mM EDTA, 20% sucrose, pH 8

*Equilibration buffer*: 20 mM imidazole in PBS, pH 8

*Wash buffer:* 10 mM NaCl, 10 mM imidazole in PBS, pH 8

*Elution buffer:* 250 mM imidazole in PBS, pH 8

*Inner cell salt buffer:* 25 mM HEPES, 110 mM KCl, 10.5 mM NaCl, 1 mM MgCl2, pH 7.5

BL21 DE3 cells (50 μL, New England Biolabs) were transformed with the pSANG10-3FBG4 plasmid^36^ (2 μL, 36 ng/μL) via heat shock: cells were thawed on ice, BG4 plasmid was added to cells and mixed gently by tapping before incubating on ice for 30 min, cells were heated to 42 ◦C for 10 sec and then left on ice for 5 min. Cells were then transferred into SOC outgrowth media (950 μL, New England BioLabs) at 37 ◦C for 1 hr before expansion into YT growth media (50 mL) containing kanamycin (50 mg/mL, Gibco) and 2% glucose. The cell suspension was grown overnight with incubation at 37 ◦C, 250 RPM. Cells from the small inoculum (25 mL) were expanded into Invitrogen Magic autoinduction media (1 L), incubated for 6 hrs, 37 ◦C, 250 RPM and then for 24 hrs, 18 ◦C, 250 RPM. Cells were pelleted by centrifugation at 4,000 g, 4 ◦C for 30 min. Cell pellets were dissolved in TES buffer (80 mL) treated with protease inhibitor tablets (Pierce protease inhibitor mini, 4 tablets). Diluted TES buffer (1:5, 120 mL), MgCl2 (2 mM) and benzonase (5 μL, Millipore) were added to the cell suspension and stirred for 30 min at 4 ◦C. Cellular lysate was then centrifuged at 8,000 g, 4 ◦C for 20 min.

The BG4 protein was purified from the cell lysate using HisPur cobalt resin (2.5 mL, Thermo Scientific). The resin was washed with Milli-Q water (12.5 mL) and equilibrated with equilibration buffer (5 mL). The equilibrated cobalt resin was then added to the supernatant of the cell lysate and stirred for 30 min. The suspension was run through a column and the resin washed twice with wash buffer (7.5 mL). BG4 was eluted by incubation of the resin with elution buffer (7.5 mL) for 15 min. The elution fraction was concentrated using a 10 kDa centrifugal filter (Amicon). The entirety of the elution was added to the column and spun down to 1.5 mL by centrifugation at 4,000 g, 4 ◦C. The protein was then dialysed by addition of inner cell salt buffer and spun down again to 1.5 mL. The concentrated protein was then aliquoted, flash frozen with liquid nitrogen and stored at -80 ◦C.

Protein purity and concentration was determined by running a 12% SDS-PAGE gel of the eluted protein alongside the Precision Plus all blue protein ladder (2 μL) and BSA standards. The gel was run at 120 V for 60 min, submerged in blocking buffer (40% ethanol and 10% acetic acid in distilled water) with shaking for 15 min and stained with coomassie blue dye (Bio-Rad) by gentle shaking overnight. Gel images were taken on an ImageQuant LAS 4000 and relative intensity of bands was determined in ImageJ.

### ChIP-seq

Intracellular salt buffer: 25 mM HEPES pH 7.6, 110 mM KCl, 10.5 mM NaCl, 1mM MgCl_2_ in Milli-Q water

Blocking buffer: Intracellular salt buffer with 1% BSA dissolved, filter sterilised before use. Wash buffer: 100 mM KCl, 0.1% TWEEN-20 and 10 mM Tris, pH 7.4, in Milli-Q water and filter sterilised before use.

ChIP-seq libraries were prepared with adaption from a previous protocol.^37^ Cells were grown to 80% confluency and then harvested at the required cell count (6 million and 10 million for PEO1 and PEO4, respectively). Cells were crosslinked in 1% formaldehyde for 10 min (50 RPM, 16 mL methanol-free formaldehyde, Pierce) and cross-linking was quenched for 10 min by the addition of glycine (1 mL, 2 M). Cross-linked cells were centrifuged for 5 min at 2000 RPM, 4 ◦C and supernatant removed. The cell pellet was then washed 3 times in ice-cold PBS, centrifuging for 5 min (2000 RPM, 4◦C) between each wash. Cell pellets were then flash frozen in liquid nitrogen and stored at -80 ◦C.

Cell pellets were thawed on ice and resuspended in ice-cold hypotonic buffer (250 μL/sample, Chromatrap) and incubated for 10 min on ice. Samples were then centrifuged for 5 min (5,000 g, 4 ◦C) and supernatant discarded. The pellet remaining was then resuspended in lysis buffer (100 μL/sample) and incubated for 10 min on ice. The suspension was sonicated for 15 cycles (30 seconds on/ 30 seconds off) at 4 ◦C (Diagenode Pico Bioruptor sonicator). Chromatin concentration was quantified using the Qubit 4 Fluorometer and Qubit dsDNA Broad Range kit. A concentration of 110-200 ng/μL was optimal. Assessment of successful chromatin sonication to a required fragment size distribution of 100-500 bp was carried out using the Agilent 4200 TapeStation system (version 4.1.1) and Agilent D5000 screentape following manufacturer’s instructions. Aliquots of chromatin were flash frozen in liquid nitrogen and stored at -80 ◦C.

Chromatin was thawed on ice before being supplemented with Triton X-100 (1%) and incubated at room temperature for 10 min. Chromatin (2.5 μL/sample), Ambion™ RNase A (1 μL/sample) and blocking buffer (45 μL/sample) was incubated together for 20 min at 1400 RPM, 37 ◦C. 400 ng of BG4 was added to the ChIP samples, and one sample was placed on ice as the input/control sample. The BG4 containing ChIP samples were incubated for 1 hr at 1400 RPM, 16 ◦C. Meanwhile, Anti-FLAG (5 μL/sample, Millipore) were washed three times in blocking buffer (50 μL/sample) with the DynaMag-2 magnet before being resuspended in blocking buffer (50 μL/sample). Anti-FLAG bead suspension (50 μL) was added to each ChIP sample and incubated for 1 hr (1400 RPM, 16 ◦C). ChIP samples then underwent three washes at 4 ◦C using ice-cold wash buffer (200 μL/sample). Two further washes were then conducted using wash buffer (200 μL/sample), with a 10 min incubation at 37 ◦C, 1400 RPM. TE buffer was then added to all samples to a final volume of 75 μL, followed by 1 μL of Proteinase K (Invitrogen, 20 mg/mL). Samples were incubated for 1 hr (1400 RPM, 37 ◦C) followed by a 2-hr incubation at 65 ◦C (1400 RPM). Samples were then purified using MinElute Reaction Cleanup Kit, following the manufacturer’s instructions, with elution in 30 μL of elution buffer before further diluting the elution to 40 μL.

qPCR analysis was conducted to ensure adequate enrichment of G4 regions over non-G4 regions prior to sequencing. Purified DNA (2 μL) was qPCR amplified using Fast SYBR Green Master Mix (10 μL) and forward and reverse primers (4 μL each, Table S4). qPCR was performed on an Agilent Technologies Stratagene Mx3005P Real-time PCR machine, with the following programme:

Cycle 1: 95 ◦C for 20 sec

Cycle 2: 95 ◦C for 3 sec

Cycle 3: 60 ◦C for 30 sec

Cycles 2-3 repeated 40 times

Percentage input values were calculated by performing the following calculation for each region: 2(Ct value of the input/Ct value for the ChIP)*100

Fold enrichment was then calculated by dividing the average percentage input of G4 regions by that of the non-G4 regions.

Libraries were prepared from input and immunoprecipitated DNA using the NEBNext Ultra II DNA Library Prep Kit for Illumina. Final libraries were subject to 100 bp paired-end sequencing at 30 million reads/sample.

### CUT&Tag

All buffers were filter sterilised before use.

*Wash buffer*: 20 mM HEPES pH 7.5, 150 mM KCl, 0.5 mM spermidine, treated with Roche complete protease inhibitor EDTA-free tablet

*Binding buffer:* 20 mM HEPES pH 7.5, 150 mM KCl, 1 mM CaCl2 and 1 mM MnCl2

*Dig wash buffer:* 0.05% digitonin in wash buffer

*Antibody buffer:* 1% BSA, 2 mM EDTA in Dig-wash buffer

Dig-300 buffer: 0.01% Digitonin, 20 mM HEPES pH 7.5, 300 mM KCl, 0.5 mM Spermidine, treated with Roche complete protease inhibitor EDTA-free tablet

*Tagmentation buffer:* 10 mM MgCl2 in Dig-300 buffer

*DNA extraction buffer:* 0.5 mg/mL proteinase K, 0.5% SDS, 10 mM Tris-HCl pH 8.

Cells were grown to 80% confluency in T75 cm3 flasks and then harvested at the required cell count (100,00 cells/sample). Cells were cross-linked in 0.1% formaldehyde for 2 min (16 mL, methanol-free formaldehyde, Pierce) and cross-linking was quenched by addition of glycine (60 μL, 1.25 M) for 5 min. Cells were then centrifuged for 4 min, 1300 g, 4 ◦C and supernatant removed. The cell pellet was re-suspended in wash buffer (100 μL/sample). Cells were next immobilised on magnetic concanavalin A (ConA) beads. ConA beads (10 μL/sample) were first washed twice in binding buffer (100 μL/sample) using a magnetic stand (DynaMag-2 magnet) and re-suspended in binding buffer (110 μL/sample). Cell suspensions (100 μL/sample) were incubated with ConA beads (10 μL/sample) for 10 min at 25 ◦C, 600 RPM. Supernatant was then removed from beads followed by two washes with wash buffer (100 μL/sample). Beads were re-suspended in antibody buffer (50 μL) with BG4 (∼5 μM, 4 μL/sample) or H3K27ac antibody (0.5 μL/sample, Millipore) and incubated at 4 ◦C overnight, 600 RPM.

Beads from BG4 reactions were washed twice with dig-wash buffer (100 μL/sample) before addition of anti-FLAG antibody (2 μL, Cell Signalling Technology) in 50 μL dig-wash buffer and incubation for 1 hr at 25 ◦C, 600 RPM. Beads were then washed three times with dig-wash buffer (500 μL/sample). All reactions were then incubated with anti-rabbit antibody (0.5 μL, antibodies-online in 50 μL dig-wash buffer) for 1 hr at 25 ◦C, 600 RPM, followed by three washes with dig-wash buffer (500 μL/sample). Epicypher pA-Tn5 enzyme was then added to each reaction (2.5 μL in 50 μL dig-300 buffer) and allowed to bind for 1 hr at 25 ◦C, 600 RPM, followed by three washes with dig-300 buffer (500 μL/sample). The tagmentation reaction was activated by addition of tagmentation buffer (300 μL/sample) for 1 hr at 37 ◦C, 600 RPM. Beads were then washed gently with TAPS buffer (Thermo Scientific), mixed with DNA extraction buffer (100 μL/sample) by vortexing for 2 sec and then incubated for 1 hr at 55 ◦C, 800 RPM. Tagmented DNA was purified using the Zymogen DNA clean and concentrator-5 kit following manufacturer’s instructions and eluted with 10 mM Tris-HCl (pH 7, 25 μL, 10 min). Purified DNA (21 μL) was PCR amplified using NEBNext HiFi PCR master mix (25 μL) and Illumina sequencing primers (2 μL universal i5 primer and 2 μL barcoded i7 primer, 10 μM, Table S5). PCR was performed on a Bio-Rad C1000 touch thermal cycler with the following programme: Cycle 1: 72 ◦C for 5 min

Cycle 2: 98 ◦C for 30 sec

Cycle 3: 98 ◦C for 10 sec

Cycle 4: 63 ◦C for 10 sec

Cycles 3-4 repeated 11 times

72 ◦C for 1 min and then hold at 8 ◦C

Amplified DNA was then purified by addition of Ampure XP paramagnetic beads (65 μL, Agencourt) and incubation at room temperature for 10 min. Beads were washed twice with 80% ethanol and DNA eluted with Tris-HCl (10 mM, 25 μL, 10 min). Fragment size distribution was analysed on an Agilent 4200 TapeStation system using the Agilent high sensitivity D1000 screentape following manufacture’s instructions. DNA concentration for each sample was quantified using the Agilent TapeStation analysis software (version 4.1.1). Samples were then pooled together so each sample was present in the pool at equivalent concentration. Large fragments (> 2,000 bp) were removed from pooled libraries by incubation of DNA with Ampure beads (0.4x volume of pool) for 15 min. Supernatant was then isolated and incubated again with Ampure beads (1.4x volume of pool) for 15 min to remove short fragments. Beads were washed twice with 80% ethanol and DNA eluted with Tris-HCl (10 mM, 50 μL, 10 min). Final libraries were subject to 150 bp paired-end sequencing at 10 million reads/sample.

### ATAC-seq

ATAC-sequencing was performed as previously described.^38^ To note, a range of 3-5 additional PCR cycles were performed per sample, dependent on the results of the qPCR, to reduce PCR bias. The customized Nextera PCR Primer sequences used are reported in Table S6. For library quantification, fragment size distribution and DNA concentration was quantified using an Agilent 4200 TapeStation system using the Agilent high sensitivity D1000 screentape following manufacture’s instructions. DNA concentration for each sample was quantified using the Agilent TapeStation analysis software (version 4.1.1). Final libraries were subject to 100 bp paired-end sequencing at 50 million reads per sample.

### RNA-seq

RNA was extracted from PEO1 and PEO4 cells (3.5 x10^6^) using Qiagen’s RNeasy extraction kit, following the manufacturer’s instructions. Briefly: adherent cells were trypsinized, counted and aliquoted. Cell pellets were then lysed with the RNeasy lysis buffer and ran through Qiagen’s QIAshedder homogenizing columns. Cell lysate was then washed with RNeasy wash buffer and an on-column DNase I digestion was performed, followed by RNA purification according to the kit’s instructions. RNA was eluted with RNase-free water (50 μL). RNA quality was analysed on an Agilent TapeStation using the RNA ScreenTape following manufacture’s instructions. RNA was sequenced via polyA enrichment and 150 bp paired-end sequencing with 20 million reads/sample.

### CUT&Tag, ChIP and ATAC-seq data analysis

Quality of reads was assessed with fastqc (http://www.bioinformatics.babraham.ac.uk/projects/fastqc/) and adapter sequences trimmed with fastp.^91^ Reads were aligned to the hg38 human genome build with Bowtie2.^92^ Reads were then filtered with samtools^93^ to remove unmapped, mitochondrial and secondary reads, black-listed regions were removed with bedtools intersect^94^ and duplicated reads removed with picard Markduplicates (http://broadinstitute.github.io/picard/) function. Peaks were called using MACS2 (–nolambda setting for CUT&TAG and ATAC-seq, default for ChIP-seq)^95^. Peaks common between replicates and peaks distinct between drug-sensitive and drug-resistant cells were identified with bedtools intersect. Genomic locations of peaks were annotated with Homer annotatePeaks^96^ and linked to the expression of the gene with the nearest TSS. Genomic tracks were visualised on IGV^97^and super-enhancers were called with ROSE.^62,63^ Motif enrichment was performed using the MEME suite online tool. Pathway enrichment was conducted with ShinyGO,^46^ GSEA (https://www.gseamsigdb.org/gsea/msigdb/human/annotate.jsp)^52^ and NDEx IQuery pathway figures tool(https://www.ndexbio.org/iquery/).^53^

### RNA-seq data analysis

Quality of reads was assessed and adapter sequences trimmed as above. Reads were aligned to the hg38 human genome build with STAR.^98^ Differential gene expression was assessed with DESeq2.^99^ Downstream analysis including sample clustering and principle component analysis were performed with iDEP.^71^Pathway enrichment was performed as above.

## Cytotoxicity assays

Sensitivity of PEO1 and PEO4 to cisplatin was measured by performing MTS assays. Cells were plated in a 96-well plate with 3 biological repeats for PEO1 and PEO4 and allowed to adhere for 24 hrs. A serial dilution (1:2, starting at 25 μM) of cisplatin (Cambridge Bioscience) was then performed into the wells. PDS was synthesised as previously described^74^ and added for synergy experiments and cells were left to grow for 3 days. Media was removed from each well and replaced with the MTS reagent (100 uL, Promega CellTiter AQueous One solution), which was allowed to develop for 2 hrs before absorbance of the plate was read at 490 nm using a CLARIOstar plus microplate reader. The percentage survival for each well was determined by calculating the relative absorbance in the control wells (no cisplatin) with that in the test wells. Curves were fit in GraphPad Prism using the Inhibitor vs. response - Variable slope model.

## Data availability

All new datasets have been deposited in the gene expression omnibus (GEO): GSEXXXXXX Previously deposited datasets used in this study: GSE149147 (ATAC-seq and RNA-seq),^24^ GSE110582 (G4-seq).^39^

## Acknowledgments

M.D.A is supported by a Biotechnology and Biological Sciences Research Council (BBSRC) David Phillips Fellowship (BB/R011605/1), a Lister Institute Research Prize and an Ovarian Cancer Action programme grant. J.R. is funded by the Engineering and Physical Sciences Research Council [EPSRC, EP/S023518/1] and the NIHR Imperial Biomedical Research Centre. G.F. is supported by the EPRSC [EP/S023518/1] and by a CRUK non-clinical training award [CANTAC721\100021]. S.G. is supported by the BBSCR [BB/W016710/1]. T.E.M. is supported by the EPSRC [EP/S023518/1]. IM is supported by an NIHR Senior Investigator award (NF-SI-0514-10101) and also acknowledges support from the NIHR Imperial Biomedical Research Centre and the Imperial Experimental Cancer Medicine Centre.

## Authors Contribution

J.R., G.F. and M.D.A. designed research with critical input from R.V., M.K., I.M., R.B. and H.K. J.R. and G.F. performed all the experiments, with support from I.G. and S.G. I.G. generated ATAC-Seq maps for PEO1^BRCA+^ and PEO4 cells. T.E.M. synthesised the PDS batch used for the synergy experiments. J.R. and M.D.A. wrote the manuscript aided by G.F. and with input from all other authors. All authors contributed to critical discussion and data interpretation. M.D.A. supervised the research.

