## Supplementary Information for "G-quadruplex structures regulate long-range transcriptional reprogramming to promote drug resistance in ovarian cancer"

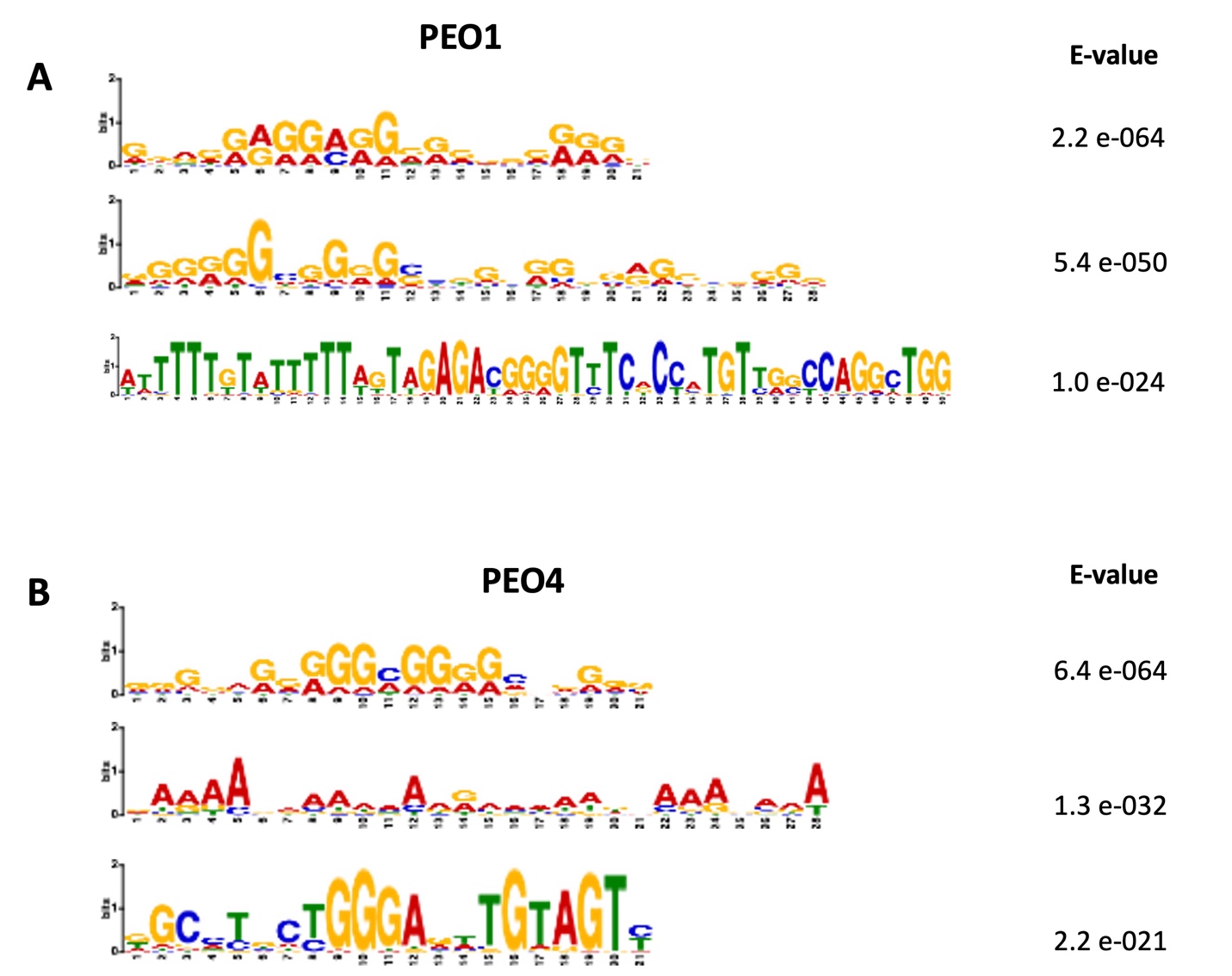

**Figure S1** – Top 3 motifs enriched amongst top 1,000 BG4 ChIP-seq peaks in A) PEO1 and B) PEO4. Motifs discovered with MEME.

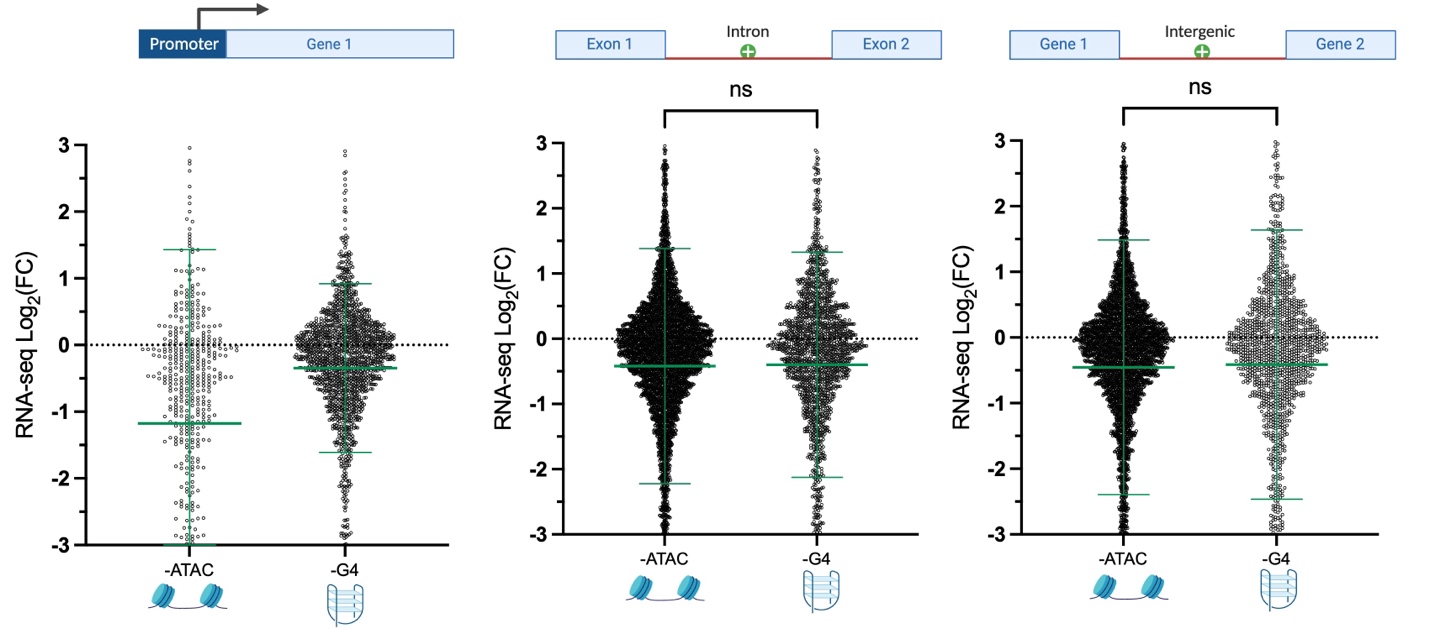

**Figure S2** – Change in expression of genes that lose an ATAC or G4 peak in PEO4 (BRCA+) relative to PEO1 (BRCA-), at promoters, intergenic and intronic regions. Only ATAC-peaks that overlap putative G4 sequences (identified for G4-seq) were examined. Statistical significance assessed by Mann-Whitney U-test. FC= fold-change; ns = non-significant; **** = p<0.0001.

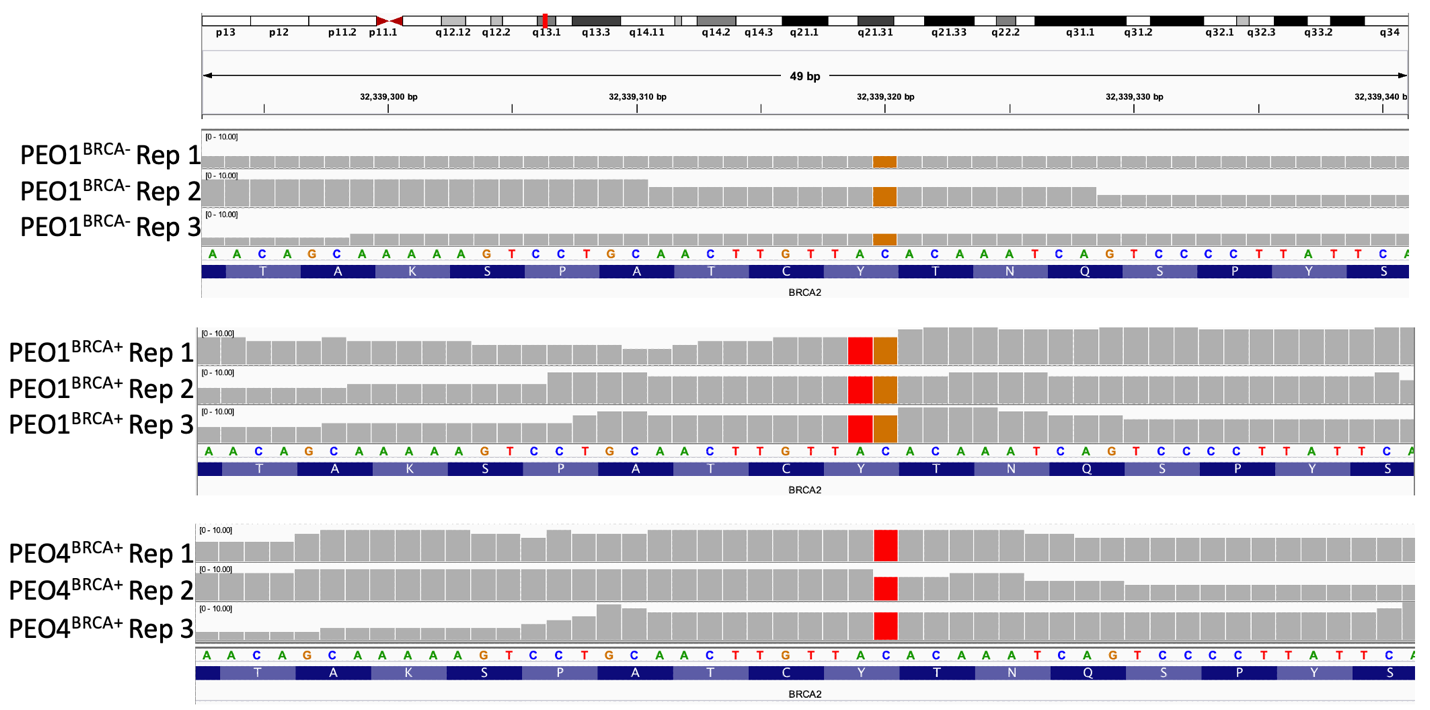

**Figure S3** – IGV profiles of RNA-seq data in PEO1^BRCA-^, PEO1^BRCA+^ and PEO4^BRCA+^ cells, showing exon 11 of BRCA2 (hg38 – chr13:32,339,293-32,339,340). PEO1^BRCA-^ harbor a single C>G point mutation, PEO1^BRCA+^ contain an additional A>T mutation and PEO4^BRCA+^ a neutral C>T alteration. Data from three biological replicates is shown.

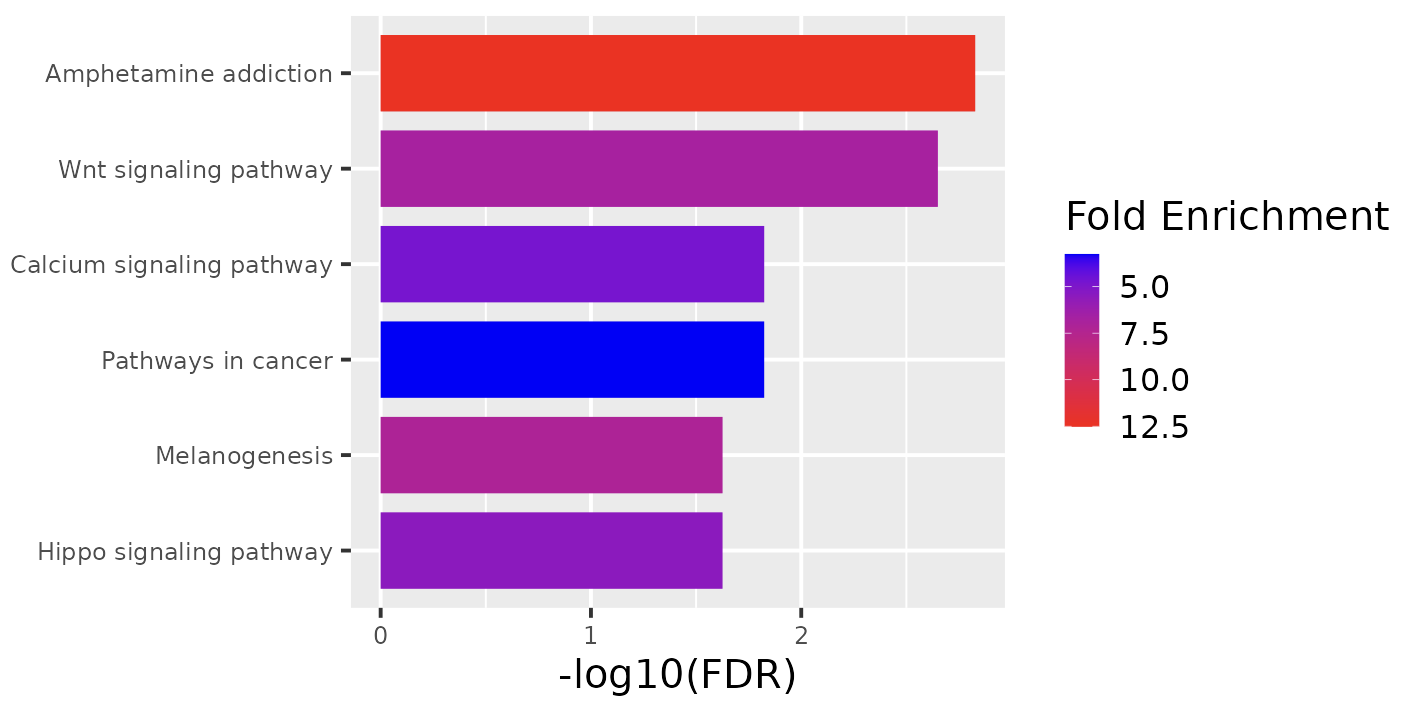

**Figure S4**– Enriched KEGG pathways of upregulated genes associated with new intergenic/intronic G4s in PEO4 (relative to PEO1^BRCA+^). Enrichment analysis performed with ShinyGO.

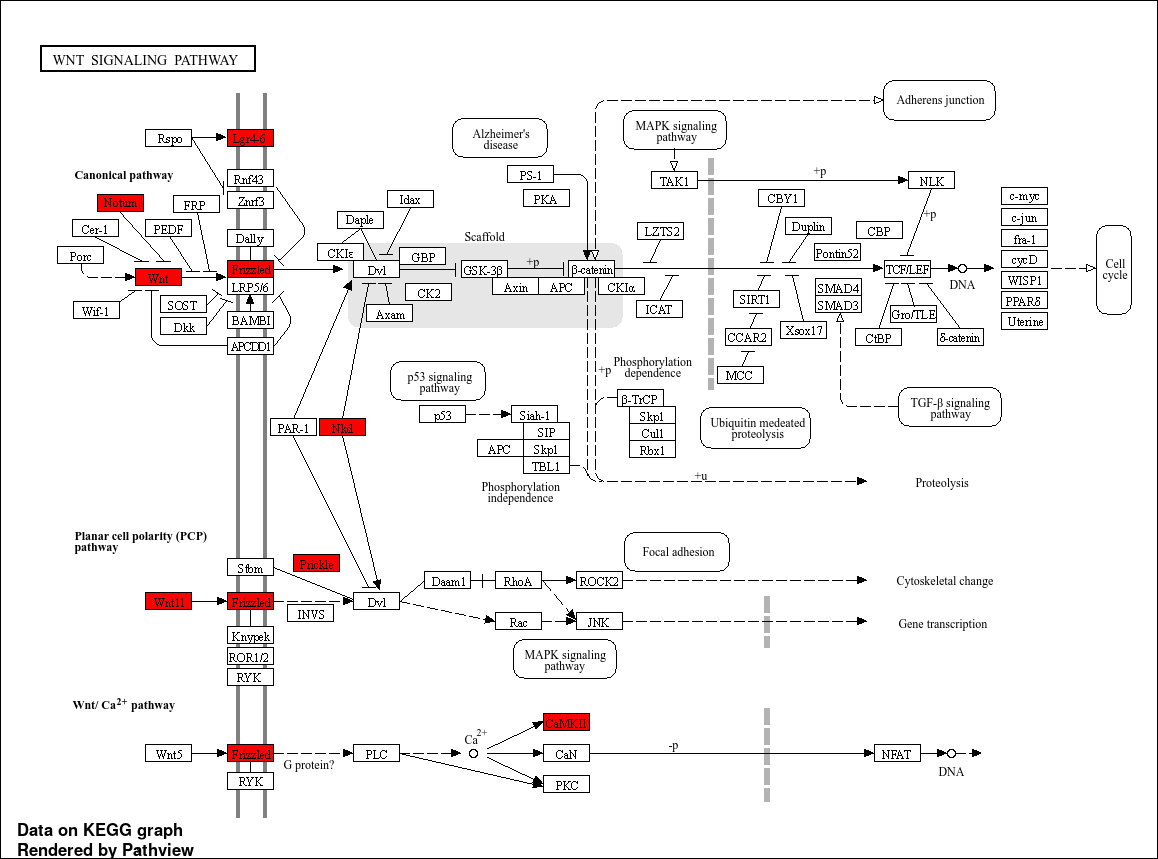

**Figure S5** – Proteins involved in WNT signaling pathway. Proteins in red are significantly upregulated in PEO4 relative to PEO1 and are proximal to PEO4-specific intergenic/intronic G4 peaks.

**Table S1** - Enriched hallmarks associated with upregulated genes proximal to new intergenic/intronic G4s in PEO4. Enrichment analysis performed with GSEA.

| **Pathway** | **FDR** |
| --- | --- |
| Estrogen response late | 1.0 e^-8^ |
| Estrogen response early | 9.5 e^-7^ |
| KRAS signaling down | 7.9 e^-5^ |
| TNGA signaling via NFKB | 5.6 e^-4^ |
| Myogenesis | 3.6 e^-3^ |
| UV response UP | 7.0 e^-3^ |
| WNT-Beta catenin signaling | 7.0 e^-3^ |
| Epithelial mesenchymal transition | 1.6 e^-2^ |

**Table S2** – Top 3 most similar publications to upregulated genes proximal to new intergenic/intronic G4s in PEO4, identified with NDEx iquery, pathway figures.

**Publication**

DNA methylation and Transcriptome Changes Associated with Cisplatin Resistance in Ovarian Cancer, Lund et al, 2017

Methylome-wide Sequencing Detects DNA Hypermethylation Distinguishing Indolent from Aggressive Prostate Cancer, Bhasin et al., 2015

The clinical applications of the cancer genome atlas project for bladder cancer, Creighton et al., 2018

**Table S3** - Top 3 most similar publications to upregulated genes proximal to new promoter G4s in PEO4, identified with NDEx iquery, pathway figures.

**Publication**

Identification of Targets of CUG-BP, Elav-Like Family Member 1 (CELF1) Regulation in Embryonic Heart Muscle, Blech-Hermoni et al., 2016

Differential Effects of Human SP-A1 and SP-A2 on the BAL Proteome and Signaling Pathways in Response to Klebsiella pneumoniae and Ozone Exposure, Wang et al., 2019

Identification of Important Effector Proteins in the FOXJ1 Transcriptional Network Associated With Ciliogenesis and Ciliary Function, Mukherjee et al., 2019

**Table S4**  – Enriched hallmarks for top 500 genes significantly upregulated in PEA2 (relative to PEA1) associated with new ATAC peaks that contain putative G4 sequences (as defined by G4-seq). Enrichment analysis performed with GSEA.

| **Pathway** | **FDR** |
| --- | --- |
| KRAS signaling up | 3.65 e-20 |
| Inflammatory response | 8.29 e-14 |
| Estrogen response early | 4.50 e-13 |
| TNFA signaling via NFKB | 4.50 e-13 |
| Epithelial to mesenchymal transition | 2.74 e-10 |
| Estrogen response late | 2.74 e-10 |
| Apoptosis | 6.64 e-10 |
| Apical junction | 1.78 e-9 |
| Complement | 1.28 e-8 |
| Allograft rejection | 6.14 e-7 |

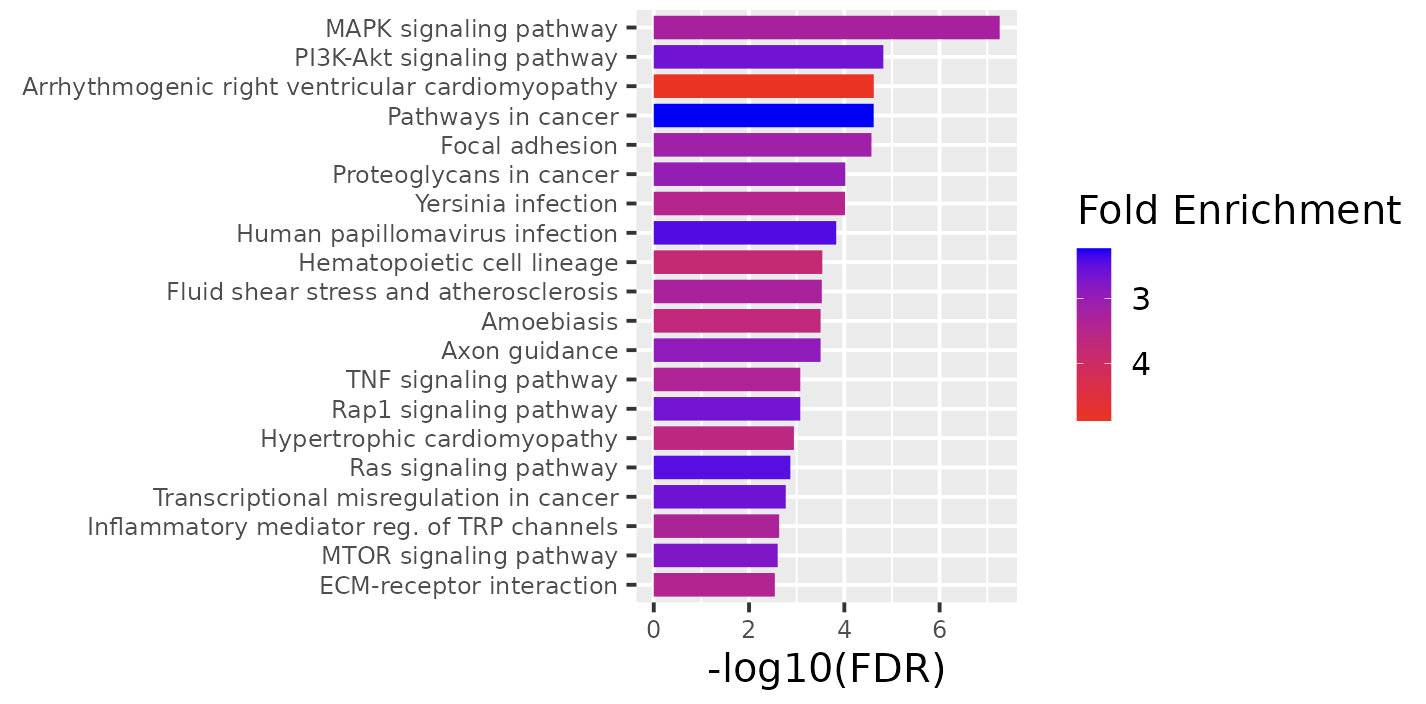

**Figure S6** – Enriched KEGG pathways of significantly upregulated genes in PEA2 (relative to PEA1), associated with new ATAC peaks that contain putative G4 sequences (as defined by G4-seq). Enrichment analysis performed with ShinyGO.

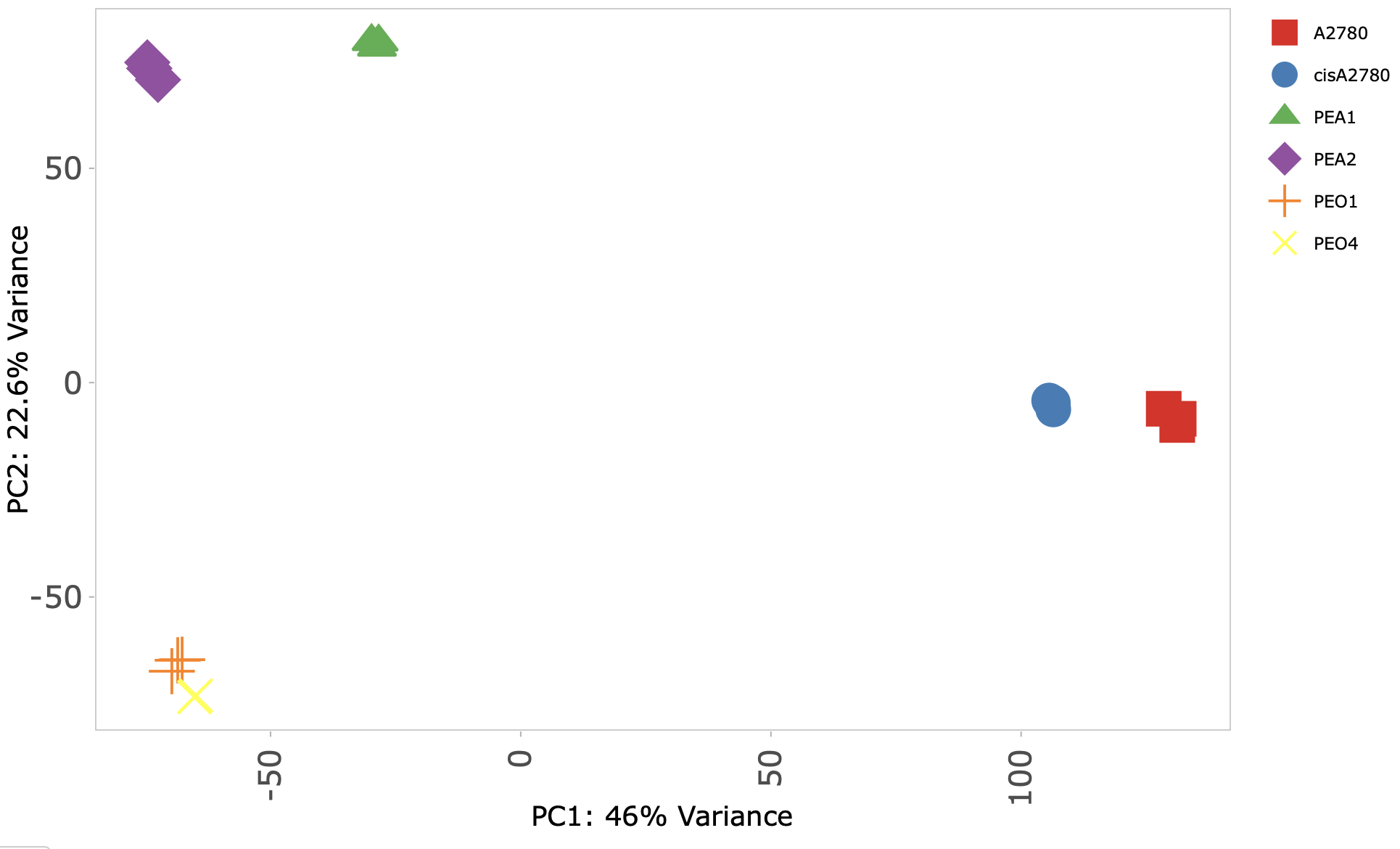

**Figure S7** – Principal component analysis of RNA-seq data in PEO1, PEO4, PEA1, PEA2, A2780 and cisA2780. Analysis performed in iDEP.

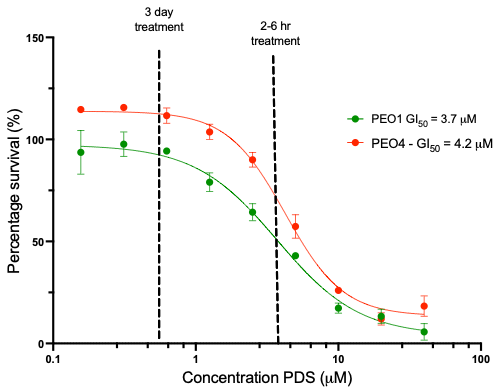

**Figure S8**– MTS viability assay of PEO1 (green) and PEO4 (red) in response to increasing concentrations of PDS, incubated for 72 hours. Error bars are standard deviations for experiments performed in triplicate.

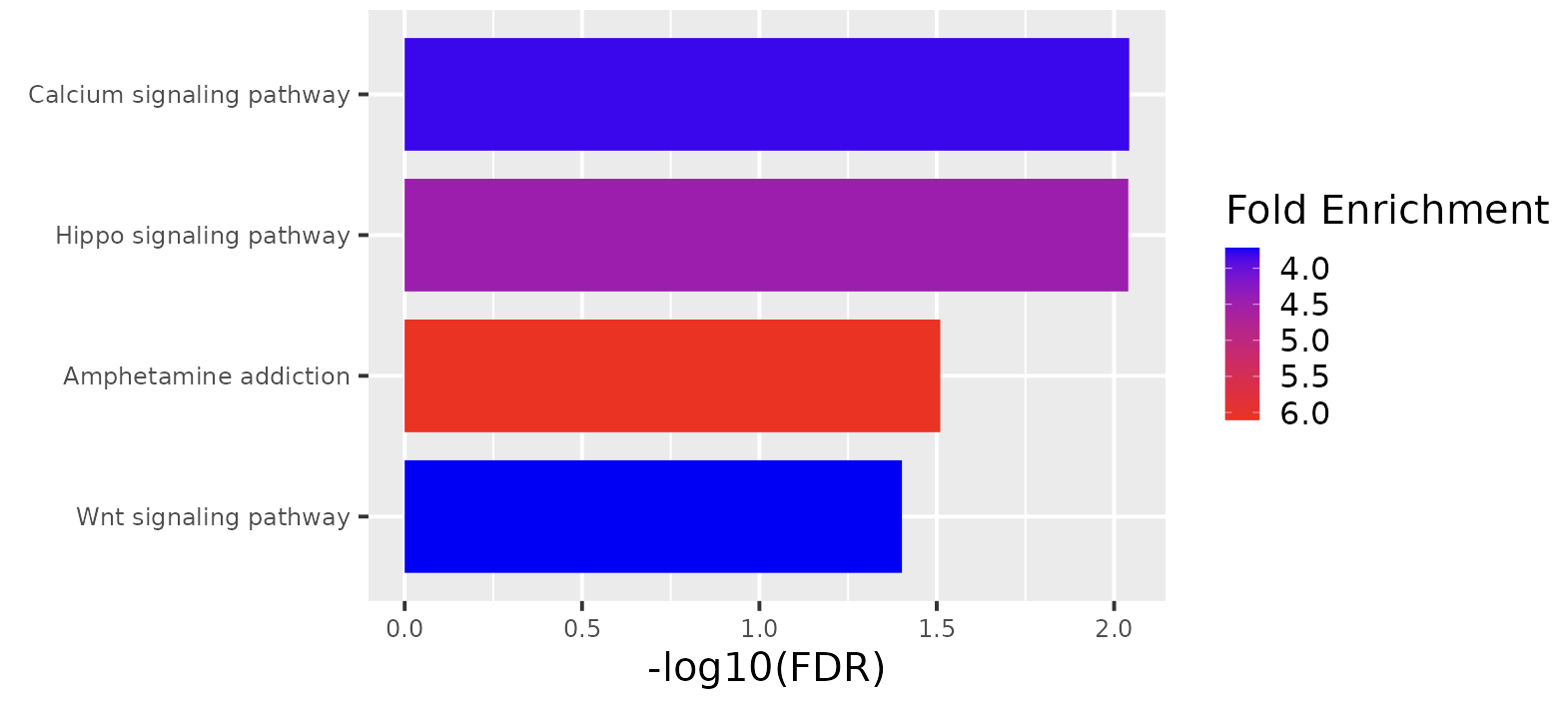

**Figure S9** – KEGG pathway enriched for genes containing intergenic/intronic G4s (detected by BG4 CUT&Tag) significantly down-regulated after 6 hours PDS treatment in PEO4. Enrichment analysis performed with ShinyGO.

**Table S4**

| **Primer region** | **Sequence (5’-3’)** |
| --- | --- |
| *MAZ* forward (G4 region) | ACT CAG CGC AGG ATT GTA AAT A |
| *MAZ* reverse (G4 region) | CCT CAT GCT TCG GCT TCC |
| *RPA3* forward (G4 region) | CGG AAG TTG ACA GAT ACA GGG |
| *RPA3* reverse (G4 region) | GAT CGC AGA AAG GTA GTC TCA G |
| *KIF14* forward (G4 region) | CGG TAG CCG TCT CTG AAT G |
| *KIF14* reverse (G4 region) | CTT TAG CAG AAC CCG AGG AG |
| *SPRED2* forward (G4 region) | AAC AGG AGG AGG AAG TAG GG |
| *SPRED2* reverse (G4 region) | TTT CGG TCG CAA GTA GGA AG |
| *TMCC1* forward (non-G4 region) | GTG GTA CAC TGC CTA CAG TAT T |
| *TMCC1* reverse (non-G4 region) | GTA TAA CGC CTG GGC TAT GT |
| *IL36G* forward (non-G4 region) | GCC CAC CTC TTT ACT TCC TTA |
| *IL36G* reverse (non-G4 region) | AAC ACT CTT TCA GCT CCA TCC |

**Table S6**

| **Primer name** | **Primer sequence (5’-3’)** |
| --- | --- |
| Ad1_noMX (Primer 1) | AATGATACGGCGACCACCGAGATCTACACTCGTCGGCAGCGTCAGATGTG |
| Ad2.1_TAAGGCGA (Primer 2) | CAAGCAGAAGACGGCATACGAGATTCGCCTTAGTCTCGTGGGCTCGGAGATGT |
| Ad2.2_CGTACTAG (Primer 2) | CAAGCAGAAGACGGCATACGAGATCTAGTACGGTCTCGTGGGCTCGGAGATGT |
| Ad2.3_AGGCAGAA (Primer 2) | CAAGCAGAAGACGGCATACGAGATTTCTGCCTGTCTCGTGGGCTCGGAGATGT |
| Ad2.4_TCCTGAGC (Primer 2) | CAAGCAGAAGACGGCATACGAGATGCTCAGGAGTCTCGTGGGCTCGGAGATGT |
| Ad2.6_TAGGCATG (Primer 2) | CAAGCAGAAGACGGCATACGAGATCATGCCTAGTCTCGTGGGCTCGGAGATGT |
| Ad2.9_GCTACGCT (Primer 2) | CAAGCAGAAGACGGCATACGAGATAGCGTAGCGTCTCGTGGGCTCGGAGATGT |
| Ad2.11_AAGAGGCA (Primer 2) | CAAGCAGAAGACGGCATACGAGATTGCCTCTTGTCTCGTGGGCTCGGAGATGT |
| Ad2.12_GTAGAGGA (Primer 2) | CAAGCAGAAGACGGCATACGAGATTCCTCTACGTCTCGTGGGCTCGGAGATGT |
